# Rescue of Conformational Dynamics in Enzyme Catalysis by Directed Evolution

**DOI:** 10.1101/185009

**Authors:** Renee Otten, Lin Liu, Lillian R. Kenner, Michael W. Clarkson, David Mavor, Dan S. Tawfik, Dorothee Kern, James S. Fraser

## Abstract

Rational design and directed evolution have proved to be successful approaches to increase catalytic efficiencies of both natural and artificial enzymes^1-3^. However, a comprehensive understanding of how evolution shapes the energy landscape of catalysis remains a fundamental challenge. Protein dynamics is widely recognized as important, but due to the inherent flexibility of biological macromolecules it is often difficult to distinguish which conformational changes are directly related to function. Here, we used directed evolution on an impaired mutant of the human proline isomerase cyclophilin A (CypA) and identify two second-shell mutations that partially restore its catalytic activity. We show both kinetically, using NMR spectroscopy, and structurally, by room-temperature X-ray crystallography, how local perturbations propagate through a large allosteric network to facilitate conformational dynamics. The increased catalysis selected for in the evolutionary screen could be rationalized entirely by accelerated interconversion between the two catalytically essential conformational sub-states, which are both captured in the high-resolution X-ray ensembles at room temperature. Our data provide a glimpse of the evolutionary trajectory of an enzyme’s energy landscape and shows how subtle changes can fine-tune its function.

---

The importance of protein dynamics in enzyme function has been under extensive investigation by experimental and computational methods and has become more widely accepted^4-8^. However, because proteins are inherently flexible, assigning a direct functional role to specific conformational changes has proved challenging. For human peptidyl-prolyl cis/trans isomerase CypA, a combination of biophysical experimental techniques has elucidated general principles of the energy landscape during catalysis^9-11^. Since evolutionary selection acts on function, a new challenge is to understand how evolution shapes these energy landscapes^12^. This challenge is best exemplified by the common implication of protein dynamics as speculative explanation for the impressive functional improvements achieved via directed evolution where often only minimal structural changes are observed^1,13^. Here, we experimentally characterize changes in the energy landscape that emerge from directed evolution of CypA for enhanced catalytic activity. We find a direct correspondence of increased protein dynamics and faster catalysis along an evolutionary trajectory.

To directly observe the changes in an enzyme’s energy landscape upon directed evolution, we turned to a previously designed second-shell mutation, S99T, in CypA that had three effects: inverting the equilibrium between two states that are essential for catalysis, decreasing their interconversion rate, and causing a parallel reduction in catalysis^11^. Can we restore the catalytic function via directed evolution and discern how the acquired mutations compensate for the impaired conformational dynamics of the S99T mutant at the molecular level? To enable directed evolution on S99T CypA, first a 96-well plate screen was developed that reports on the enzymatic activity of CypA. Proline isomerase activity is difficult to screen because of its high thermal background rate (2–9 × 10^−3^ s^−1^)^14,15^. Additionally, there are several proline isomerases in *Escherichia coli*, which complicates screening in cell lysate. To overcome these limitations, we took advantage of the *Pseudomonas syringae* phytopathogenic protease AvrRpt2, which is activated by eukaryotic, but not prokaryotic, cyclophilin homologs^16^. We expressed a library of CypA S99T variants created by random mutagenesis, added inactive AvrRpt2 to cell lysate and monitored the cleavage of an AvrRpt2 substrate^17^ (Fig. 1a). Besides revertants to wild-type CypA (Ser99), we identified a variant (S99T/C115S) with increased activity (Fig. 1b). A second round of screening of S99T/C115S CypA identified an additional mutation (I97V) with a further increase in activity. Extensive efforts to further improve enzymatic activity by many more rounds of evolution were unsuccessful. Both gain-of-function mutations are in proximity of Thr99, but not in direct contact with the peptide substrate (Fig. 1c). Each mutation contributes additively to the increase in activity, measured as k_cat_/K_M_^18^, and is consistent across two substrate peptides (Fig. 1d). Substrate binding affinities are only slightly changed relative to the S99T mutant (Fig. 1e and Extended Data Fig. 2) suggesting that the two mutations function by modulating the turnover rate rather than substrate binding.

**Figure 1.**
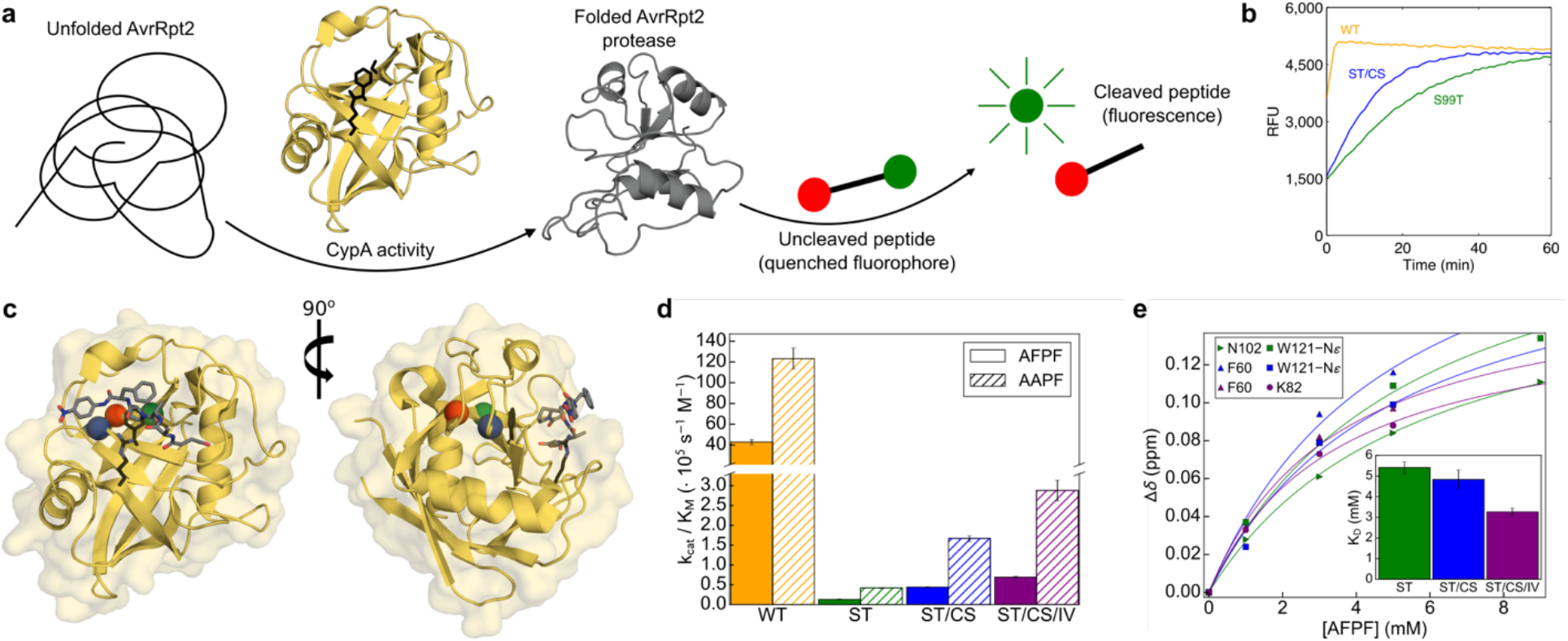
Directed Evolution selects rescue mutations for catalysis. **(a)** Scheme of the assay used in directed evolution to identify CypA mutations with increased catalytic activity: CypA activity for folding of AvrRpt2 protease measured by AvrRpt2-mediated cleavage of the peptide Abz-IEAPAFGGWy-NH2 (y = 3-nitro-Tyrosine). **(b)** Assay of directed evolution performed on cell lysate in 96-well plates to identify rescue mutations for S99T with increased CypA activity. Kinetics of peptide cleavage is shown for wild-type (yellow), S99T (green) and S99T/C115S (blue) CypA. **(c, d)** The severely catalytically compromised S99T mutant (green) is rescued by second-shell mutations (C115S, blue and I97V, red). **(c)** Sites of mutations are plotted onto the crystal structure (1RMH^29^) of CypA bound to Suc-AAPF-pNA (grey sticks) and the active-site residues R55 and F113 are shown in black stick representation. The overlay of NMR spectra shows that the overall structure of all CypA forms is very similar, with perturbations observed for residues close to the mutation site or in the dynamic network (Extended Data Fig. 1). **(d)** k_cat_/K_M_ values for wild-type, S99T, S99T/C115S and S99T/C115S/I97V CypA measured by protease coupled hydrolysis^18^ of Suc-AFPF-pNA and Suc-AAPF-pNA peptides. Error bars indicate the standard deviation obtained from triplicate measurements on at least three different enzyme concentrations. **(e)** K_D_ values for the three mutant forms of CypA for Suc-AFPF-pNA measured by NMR chemical shift analysis from peptide titrations (see also Extended Data Fig. 2). Error bars denote the standard errors in the fitted parameters obtained from the global fit.

To determine whether the rescue in catalysis is due to faster protein motion, we performed NMR dynamics experiments on the S99T, S99T/C115S and S99T/C115S/I97V mutants (Fig. 2 and Supplementary Data 1-6). Previous NMR CPMG dispersion experiments suggested a direct link between the speed of a conformational change in a dynamic network, labeled group-I, for both wild-type (WT) and S99T CypA (Fig. 2a, red)^10,11^. In contrast, a second dynamic process, group-II, comprised of loops adjacent to the active site (Fig. 2a, blue), is insensitive to mutation and displayed faster dynamics. Despite the lack of correlation between the dynamics and catalysis in the group-II residues in S99T, these residues have recently been proposed to be directly linked to catalysis in CypA^19^. To understand the changes in the energy landscape during the directed evolution that led to faster enzymes, we first turned to well-established ^15^N-CMPG experiments. Interestingly, the exchange contribution, R_ex_, for group-I residues gets bigger with consecutive rescue mutations (Fig. 2b), indicating that these mutations increase the slow interconversion rate from the major to minor state (i.e., R_ex_ = 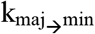). We note that the fast loop motion of group-II (residues 65-80) is observed in all enzyme forms and remains essentially unaltered (Fig. 2a-c,e,g and Supplementary Data 1-3).

**Figure 2.**
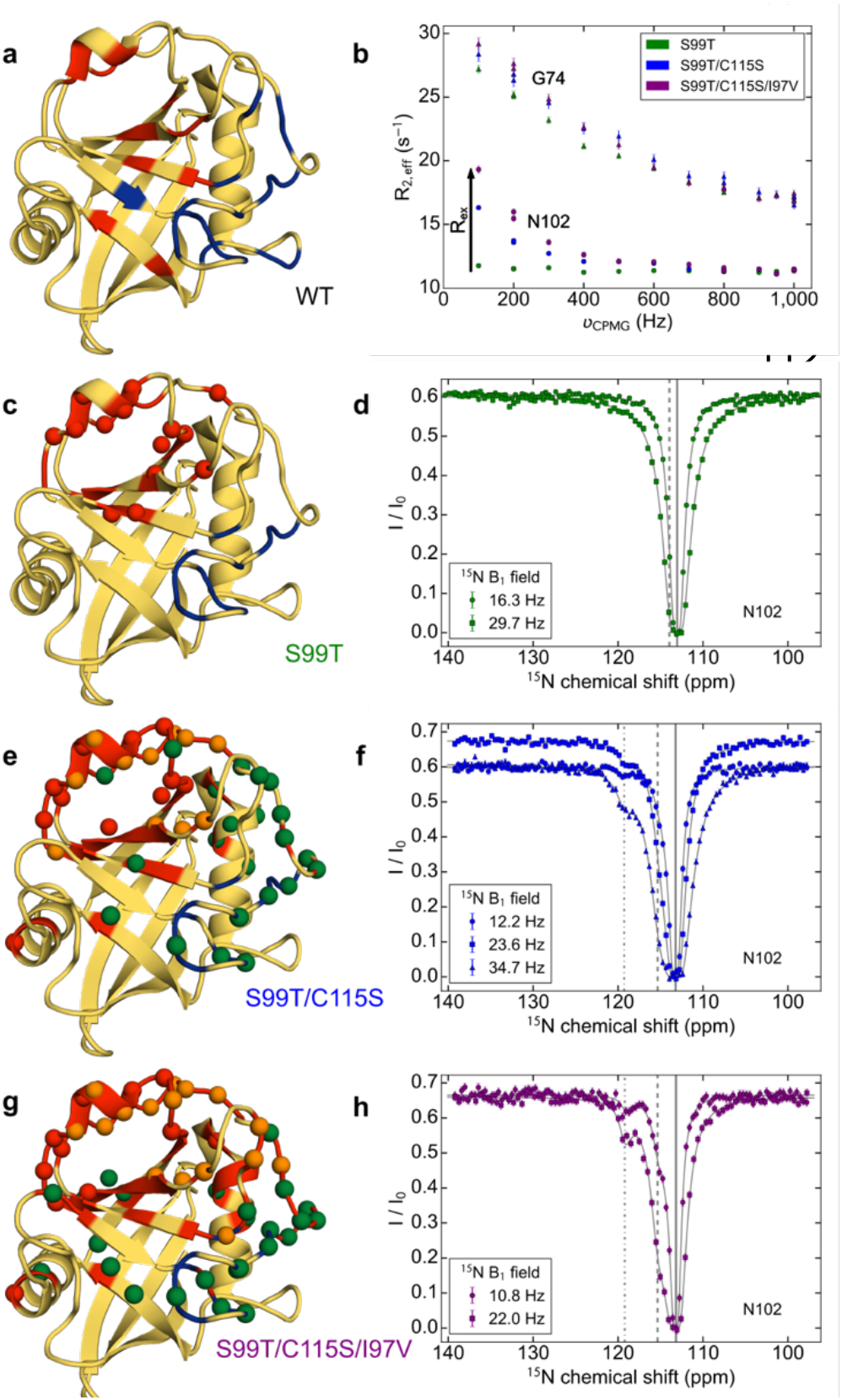
Rescue mutants alter the conformational dynamics of CypA as measured by NMR. **(a)** Dynamics on WT CypA (shown here)^10^ and S99T^11^ identified a slower (group-I, red) and faster dynamical process (group-II, blue). **(b)** For the three mutants, ^15^N CPMG dispersion profiles for a representative residue in fast exchange (Gly74) from group II and slow exchange (Asn102) from group I^11^. The fast-exchange process is virtually unaltered by the mutations, whereas Rex increases consecutively from single via double to triple mutant (see also Supplementary Data 3). **(c, e, g)** Quantitative analysis of fast and slow protein dynamics of CypA rescue mutants by CPMG relaxation and CEST experiments are plotted onto the structure. Blue and red color coding of the cartoon representation denotes fast and slow dynamics as determined from the temperature-dependence and shape of CPMG relaxation dispersion profiles (Supplementary Data 1 and 2). Spheres represent residues in slow exchange with quantifiable CEST profiles. **(c)** CEST data for all 15 residues (red spheres) in S99T can be globally fit to a two-site exchange process (Extended Data Fig. 3 and Supplementary Data 4). **(e, g)** Residues with CEST profiles in S99T/C115S **(e)** and S99T/C115S/I97V **(g)** are well-described by two distinct slow processes (red and green, respectively), whereas residues shown in orange sense both processes and require a three-site exchange model (Extended Data Figs. 4 and 5 and Supplementary Data 5 and 6). **(d, f, h)** Representative ^15^N CEST profiles of residue N102, measured at the indicated field strengths are shown for single **(d)**, double **(f)**, and triple **(h)** mutants of CypA. The chemical shifts for the major (solid line) and minor states (– – – and – · – lines) are indicated. Uncertainties in R_2,eff_ (panel b) are determined from the rmsd in the intensities of the duplicate points (n = 4) according to the definition of pooled relative standard deviation; uncertainties in I/I0 for CEST data (panels d, f, h) are estimated from the scatter in the baseline of the profile (typically, n > 50).

For a quantitative understanding of the mechanism underlying the increased catalysis along the directed evolution trajectory, we applied a powerful NMR method for studying systems in slow exchange, chemical exchange saturation transfer (CEST) spectroscopy^20^. The ^15^N-CEST experiments identified a large number of residues in slow exchange in all mutant forms of CypA (Fig. 2c-h; Extended Data Figs. 3-5 and Supplementary Data 4-6) and delivered two key results. First, for the rescue mutants we observe two distinct slow dynamic clusters that differ in kinetics and populations (Fig. 2e,g and Extended Data Figs. 4 and 5). The exchange profiles of 46 residues for the double-, and 55 residues for the triple-mutant were globally fit to a linear, three-state exchange model (see Online Methods). Second, we discover a gradual increase in the interconversion rate from the major to the minor conformation for the slow process centered around group-I residues (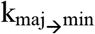 2.3 ± 0.1 s^−1^ for S99T, 8.2 ± 0.4 s^−1^ for S99T/C115S and 11.5 ± 0.5 s^−1^ for S99T/C115S/I97V, respectively).

NMR relaxation experiments enable us to “see” the prevalence of exchange on different time scales and to unravel their importance in biological processes^4^. The remarkable correspondence between 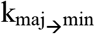 and k_cat_/K_M_ (Fig. 3a) corroborates our hypothesis that the increased dynamics of group-I residues is indeed responsible for the rescue in enzymatic activity. The good correlation between the chemical shift differences, Δδ_AB_ and Δδ_AC_, in the two rescue mutants (Fig. 3b) indicates that the exchange processes are the same by nature. The newly identified second slow exchange process is located on the opposite side of the protein and partially overlaps with the faster loop motion detected by CPMG experiments (Fig. 2e,g). The rather large chemical shift differences observed for this exchange process (Fig. 3b, green squares) agree well with the predicted values of going to a more extended/unfolded state (Extended Data Fig. 6) and are, therefore, unlikely to be relevant for catalysis.

**Figure 3.**
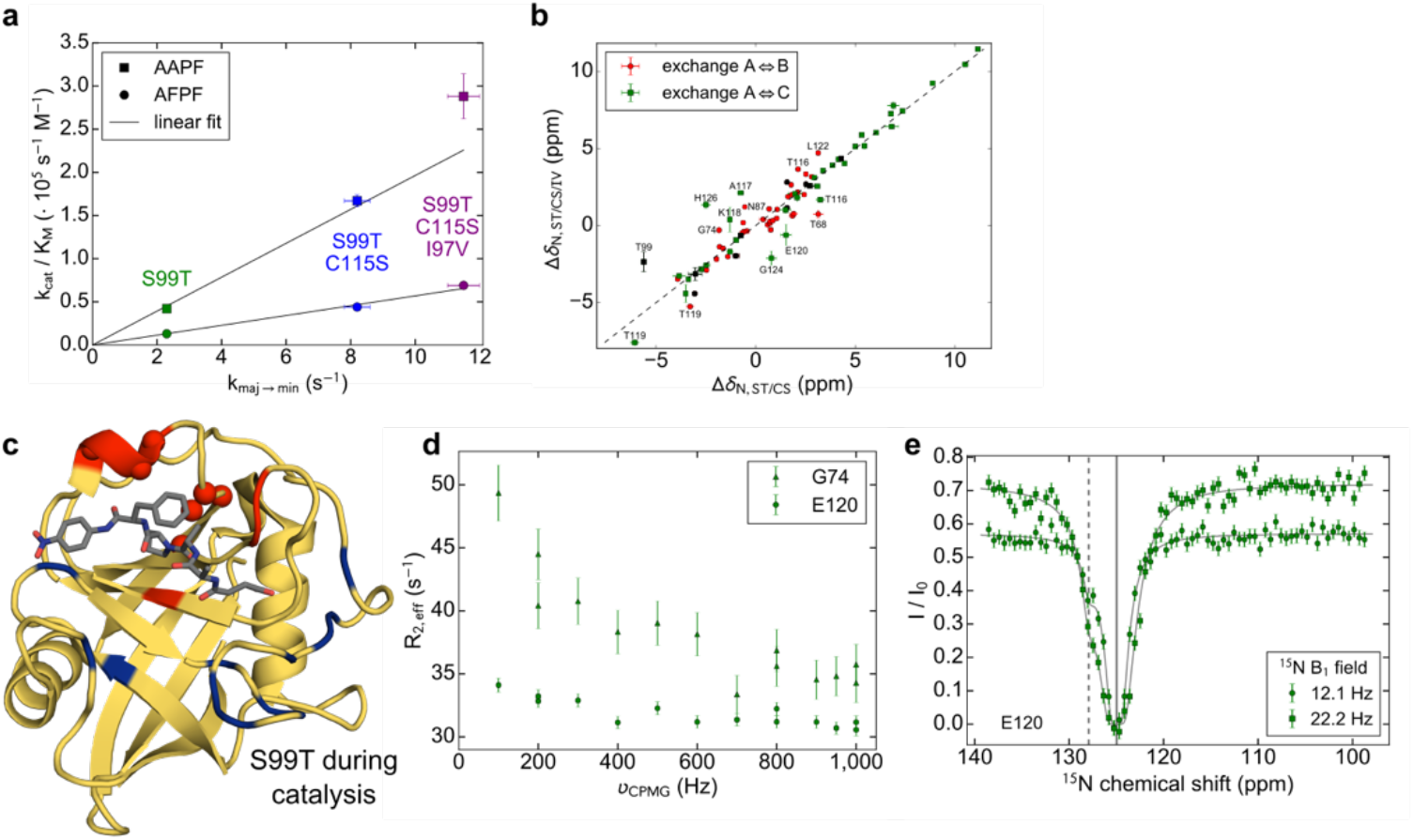
Protein dynamics during turnover and catalytic efficiency correlate. **(a)** Correlation between k_cat_/K_M_ and 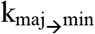 measured by CEST across all rescue mutations for apo protein. **(b)** Correlation of the chemical shift differences between major and minor conformations for the two processes observed in S99T/C115S and S99T/C115S/I97V. Residues within 5 Å of mutation site (I97V) are shown in black and assignments are given if the variation in Δδ is >1.5 ppm. **(c)** Quantitative analysis of fast and slow protein dynamics of CypA S99T during catalysis of Suc-AFPF-pNA peptide (grey sticks) measured by NMR. CPMG relaxation dispersion experiments revealed, similarly as in apo S99T, fast motion mainly in the flexible loop (blue) and a slow process (red) consistent with the CEST data (spheres). **(d)** Representative CPMG profiles for a residue in fast (Gly74) and slow (Glu120) exchange and **(e)** CEST profile for Glu120 during catalysis (see Supplementary Data 7 and 8 for all profiles). Error bars in panels a and b denote the (propagated) standard errors in the fitted parameters. The uncertainties in CPMG **(d)** and CEST **(e)** data are determined as described in Fig. 2.

Since overall turnover in WT is dictated by 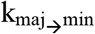 and occurs on similar timescales in both the apo enzyme and during turnover^10^, we needed to confirm that such a correspondence between the measured dynamics in the apo and turnover protein holds true for the mutants. Therefore, CPMG and CEST experiments were performed on S99T during enzymatic turnover of a substrate peptide. These experiments on the mutants proved to be difficult and only possible for S99T due to stability, and the weak affinity for the peptides allowed for a maximum of ~70% saturation. For S99T during catalysis, we indeed observe both fast loop movement, and slow conformational dynamics in the group-I residues, very similar to the apo protein (Fig. 3c-e). Together these data show that the intrinsic dynamics in the group-I residues in the mutants are rate-limiting for the catalytic cycle.

To reveal the structural basis of the increased dynamics in the group-I residues and hence increased catalysis along the evolutionary trajectory, we collected room-temperature X-ray data for S99T/C115S/I97V (Fig. 4). Alternate conformations were identified using qFit^21^ and the final multiconformer model was obtained after subsequent manual adjustments and occupancy refinement (Extended Data Fig. 7). A swap of the major/minor states from WT to S99T/C115S/I97V is observed for the group-I residues, thereby directly delivering the atomic structures of the conformations for which we measured their interconversion rate by NMR. Both the C115S and I97V mutations subtly reduce the amino acid size, and combined partially restore the “Phe113-in” conformation as a minor conformation that can now be directly observed in the electron density (Fig. 4b,d-f). The interpretation of increased conformational heterogeneity of Phe113 and Thr99 was additionally confirmed by an alternative ensemble refinement method (Extended Data Fig. 8). The size reduction by the C115S mutation could contribute to the faster transition between the major and minor conformation due to the relief of a clash between the larger sulfur atom and Thr99 (Fig. 4e).

**Figure 4.**
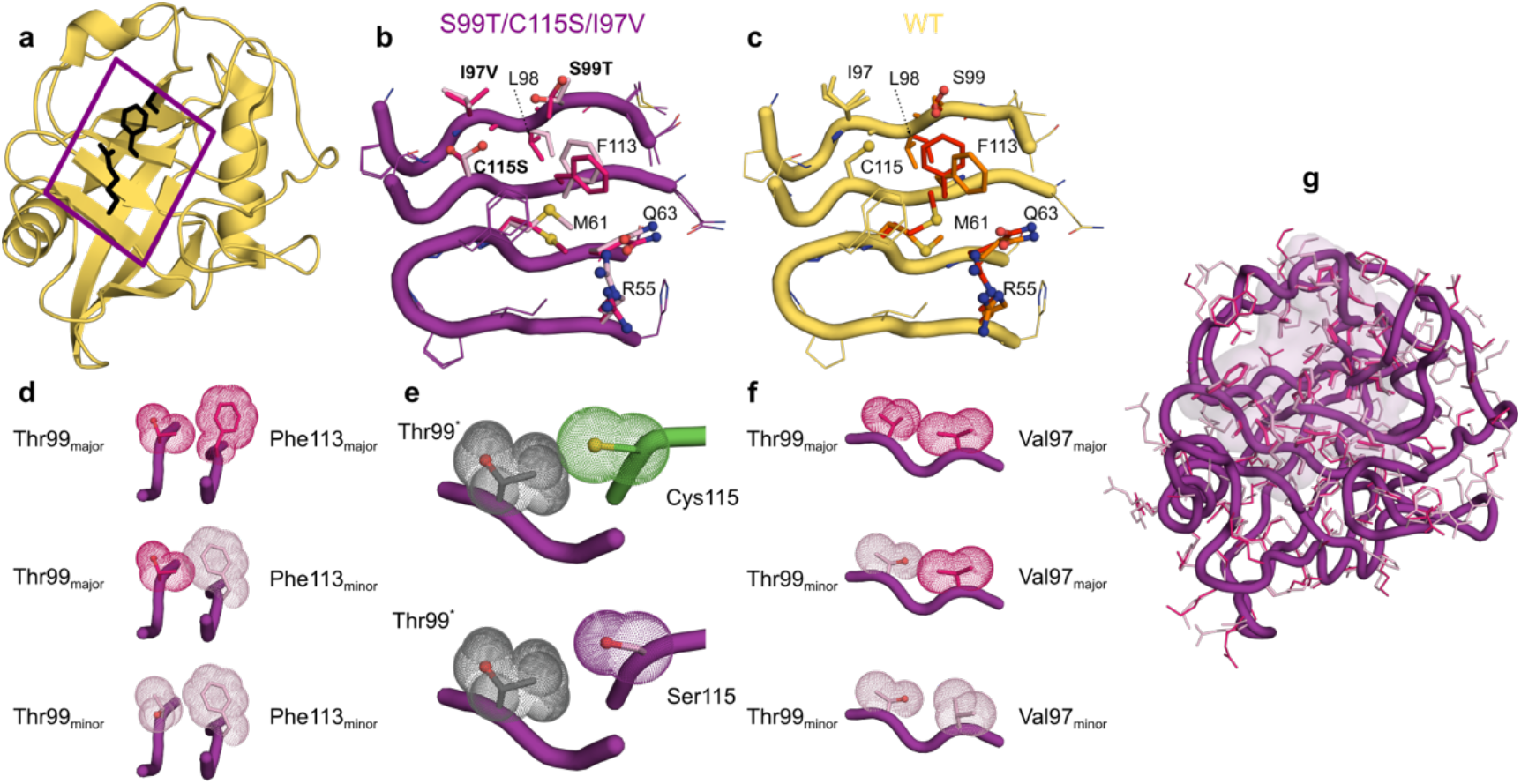
Structural basis of the increased protein dynamics from room-temperature X-ray crystallography on rescue mutants. **(a)** X-ray structure of CypA (2CPL^30^) with the active-site residues Arg55, Ser99 and Phe113 shown in black stick representation. The boxed area indicates the extended dynamic network shown in more detail in b-c. **(b)** Major and minor side chain conformations are shown for S99T/C115S/I97V (5WC7, 1.43 Å, see Extended Data Table 1) in purple and pink, respectively. The populations are flipped relative to wild-type CypA (3K0N, 1.4 Å)^11^ **(c)**, where major/minor states are shown in red and orange, respectively. **(e, f, g)** Less steric hindrance due to the reduced size of side chains in rescue mutants facilitates the interconversion between major and minor conformations. Coupling between the conformation of Thr99 and Phe113 **(e)** and Val97 and Thr99 **(g)** are necessary to relieve clashes. **(f)** The C115S (purple) mutation allows for a transition between Thr99 conformations (grey indicates morph between the major in minor state labeled Thr99*) without clash, in contrast to the bulkier Cys residue (green). **(h)** CONTACT analysis^22^ of alternative conformations of S99T/C115S/I97V identifies a network extending across group-I residues (pink surface representation) consistent with the NMR results.

The connection between these alternate conformations and the collective fitting of the NMR dynamics is further buttressed by analysis of contacts between alternative conformations^22^, which identifies a network across the protein that coincides with the group-I residues (Fig. 4g). While the detection of alternate conformations by X-ray diffraction does not per se deliver information about correlated motions, a detailed analysis of the steric constraints of group-I residues (Fig. 4d-f) exposes how motions from the active site are propagated in a correlated manner. These structural results, together with the characterization of collective dynamics by NMR, reveal how the mutations selected by directed evolution have rewired the internal packing to increase the dynamics of Phe113 correlated with surrounding group-I residues during the catalytic cycle.

Why did additional rounds of directed evolution fail to yield further improvement in the catalytic rate? We speculate that the enzyme may be in a local minimum in the fitness landscape, and that a specific combination of mutations is needed for further improvement, including mutations that are neutral or mildly deleterious mutations on their own, in agreement with the dominance of epistasis in protein evolution^23^.

Characterizing how directed evolution shaped the energy landscape for enhanced catalytic activity by solely increasing the conformational interconversion rates in a specific dynamic network has broad implications for resolving controversies about the role of protein dynamics in enzyme catalysis of modern enzymes, and in discovering the mechanism of improved catalysis via directed evolution. We briefly discuss both points in respect to pertinent current views in these fields. Fueled by a strong dispute about protein dynamics impact on catalysis^24,25^, the field has focused on characterizing several enzymes mechanistically in great detail using a combination of experimental and computational approaches, including CypA^9-11,19,26^. A two-state ensemble calculation using exact NOEs as constraints revealed an open- and closed-state of free CypA with respect to the position of the 64-74 loop^19^. The authors postulate that this loop dynamics is directly linked to catalysis via long-range concerted motions extending to the active site. This model is incompatible with the loop dynamics measured here, which remains fast for all CypA forms and is not correlated to catalytic turnover. In contrast, the group-I residues could be directly linked to catalysis via concurring changes in dynamics between WT, S99T, and the evolved enzyme forms, and catalytic turnover rates. This highlights the importance of NMR dynamics measurements and their quantitative comparison to corresponding changes in catalytic rates, which cannot be extracted from an ensemble-averaged NOE-based structure calculation.

The slowness of conformational interconversion in CypA requires enhanced computational sampling methods for MD simulations. Transitions between the experimentally determined conformational states^11^ were calculated using parallel-tempering metadynamics resulting in a free energy difference for S99T^27^ that is in excellent agreement with the experimental value of −2.5 kcal/mol obtained here from the CEST data. A two-state ensemble for the cis- and trans-peptide bound CypA calculated by replica-exchange MD simulations in combination with NMR constraints^26^ show only minimal protein conformational differences compared to the starting crystal structures (Extended Data Fig. 9). This is in sharp contrast to our NMR dynamics for WT^10^ and S99T during catalysis measured here, which clearly shows that conformational sub-states interconvert across the core catalytic network and that this rate is correlated to catalysis.

It is notable that the function of CypA was modulated by mutations that do not directly contact the substrate. While the results of numerous “second shell” mutations emerging from directed evolution experiments have been interpreted based on speculative links between protein dynamics and changes in activity, experimental evidence of alterations of populations or kinetics of alternative conformations has been sparse^13^. Here, the increased dynamics can be rationalized by the ability of Phe113, which directly abuts the substrate, to transition between different rotameric states. This transition is controlled by the repacking of alternative conformations of core residues, such as Thr99, and is enabled by the decreased bulk of the mutated residues (Cys>Ser; Ile>Val). NMR spectroscopy directly reveals how the kinetics of these transitions between alternative internal packing arrangements controls the increased catalytic activity accumulated during directed evolution. The slow dynamics associated with the catalytic cycle of CypA, and likely for many other enzymes, are only now becoming accessible to molecular simulations^28^. Therefore, the lessons of how non-active site mutations can alter conformational dynamics, derived from our integrated analysis of room-temperature X-ray data and NMR dynamics measurements, could be applied more broadly to aid the long-term goal of rationally improving directed evolution.

## Methods

### Library Screen

Mutant libraries were created using error prone PCR as in Rockah-Shmuel *et al.*^31^. For the initial library used to isolate S99T/C115S and subsequent screening efforts, the mutation rate was tuned to <3 new mutations per gene. Over 1,000 individual clones were screened in the initial screen that identified S99T/C115S and an additional 1,500 clones were screened in a second library to identify S99T/C115S/I97V. Subsequent screens of more than 5,000 variants, including random mutations and focused libraries randomizing contacting residues, to identify additional mutations did not yield any new mutations with gains of function.

From transformations of >10,000 individual colonies, individual isolates were picked into 96 well blocks and grown overnight at 37 °C. For induction, 4 μl of the overnight culture was diluted into a fresh 96 well block containing 1 mL of LB and grown for 3 hours prior to addition of 100 μM IPTG. The induced culture was then grown overnight at room temperature and harvested by centrifugation at 3000xg for 15 minutes. The media was decanted from the 96 well block and 125 μL of lysis buffer (20 mM Tris, pH 8.0, 1% Triton S100) was added. The block was shaken for 30 minutes at room temperature and then frozen at −80 °C.

Prior to assaying the AvrRpt2 activity, the 96 well block was thawed. To remove the membrane and unlysed cells, the thawed lysate was centrifuged at 3000xg for 15 minutes. 30 μL of the extract was transferred carefully to a 96 well plate. A master mix of inactive 0.25 mg/mL AvrRpt2 and 1 mM substrate peptide (Abz-IEAPAFGGWy-NH2, where y = 3-nitro-Tyrosine) were mixed in reaction buffer (20 mM HEPES, pH 8.5, 50 mM NaCl, 1 mM DTT). The reaction was initiated by mixing 10 μL of lysate with 30 μL of master mix in a Costar Black Flat bottom 96 well plate (Corning, product #3694) and measured in a Safire microplate reader monitoring fluorescence at 418 nm (excitation at 340 nm).

### Sample Preparation

Wild-type CypA and mutant proteins were essentially expressed and purified as described previously^9^. Briefly, LB medium or M9 minimal medium containing 2 g/L U-[^13^C]-D-glucose and/or 1 g/L ^15^NH4Cl (Cambridge Isotope Laboratories, Tewksbury, MA, USA) as the sole carbon and nitrogen source were used to express unlabeled and ^13^C/^15^N-labeled or ^15^N-labeled CypA, respectively. Cells were grown at 37 °C to an OD_600_ ~0.6 after which protein expression was induced with 0.3 mM IPTG for 4 hours at the same temperature (wild-type and S99T) or overnight at 20 °C (S99T/C115S and S99T/C115S/I97V). Cells were lysed in 25 mM MES, pH 6.1, 5 mM b-mercaptoethanol and purified on a SP-Sepharose column using a NaCl gradient. Fractions containing CypA were pooled and dialysed overnight into 50 mM Na_2_HPO_4_, pH 6.8, 5 mM b-mercaptoethanol. Remaining DNA and other impurities were removed using a Q-Sepharose column by collecting the flow-through. CypA was purified to homogeneity on a size-exclusion column (HiLoad 16/600, S75) equilibrated in 50 mM Na_2_HPO_4_, pH 6.5, 250 mM NaCl, 5 mM β-mercaptoethanol. Samples were dialysed overnight into their final buffers (50 mM Na_2_HPO_4_, pH 6.5, 1 mM TCEP for NMR and 20 mM HEPES, pH 7.5, 100 mM NaCl, 0.5 mM TCEP for activity measurements, respectively). All purification steps were performed at 4 °C.

### Crystallography

S99T/C115S/I97V CypA was produced and crystallized similarly to previous studies of wild-type CypA^11^. Briefly, crystals were grown by mixing equal volumes of well solution (100 mM (4-(2-hydroxyethyl)-1-piperazineethanesulfonic acid) HEPES, pH 7.5, 23% PEG 3350, 5 mM Tris (2-carboxymethyl) phosphine [TCEP]) and protein (60 mg/mL in 20 mM HEPES, pH 7.5, 100 mM NaCl, 0.5 mM TCEP) in the hanging-drop format. To collect the room-temperature synchrotron dataset, paratone oil was applied to cover a 2 μL hanging drop containing a single large crystal of S99T/C115S/I97V CypA. The crystal was harvested through the paratone and excess mother liquor was removed using a fine paper wick. Attenuated data were collected at ALS beamline 8.3.1 with a collection wavelength of 1.115 Å at 273 K controlled by the cryojet using an ADSC Q315r detector. Data were processed using XDS^32^, monitoring scaling statistics to confirm a lack of radiation damage^33^ and CC statistics for high resolution cutoffs^34^, to 1.43 Å resolution.

Molecular replacement was performed in Phaser^35^ using 2CPL^30^ as an initial search model. Residues were manually mutated in Coot^36^ and subjected to multiple rounds of refinement using phenix.refine^37^. To add alternative conformations in a systematic manner, the refined single conformer model was rebuilt using qFit^21^. To finalize the model, further manual improvements to the connectivity of alternative conformations and the ordered solvent molecules were performed iteratively with cycles of phenix.refine. Structure validation was performed using MolProbity and yielded the following statistics for Ramachandran (favored: 96%, allowed 4%, outliers: 0%), 1.4% rotamer outliers and a clashscore of 0.98.

Analysis of contacting residues using CONTACT was performed (parameters: Tstress 0.35 and Relief 0.9), as for wild-type CypA^22^. Ensemble refinements were performed using phenix.ensemble_refinement^38^. All figures were prepared using PyMol^39^.

### Enzyme activity

*k*_cat_/*K*_M_ for the enzyme catalyzed *cis*-to-*trans* isomerization of succinyl-AAPF-p-nitroanilide and succinyl-AFPF-p-nitroanilide (Suc-AXPF-pNA; Bachem) was measured at 10 °C using the standard chymotrypsin-coupled assay^18^. The increase in absorbance at 390 nm was fit to a single exponential to yield a rate constant for the interconversion between *cis*- and *trans*-peptide. Both the uncatalyzed, background (triplicates) and enzyme-catalyzed reaction were (average/standard deviation of at least three different enzyme concentrations, each measured in triplicate) measured for both peptides and different enzyme concentrations chosen such that the rate of the enzyme-catalyzed reaction is between 3- and 15-fold faster than the uncatalyzed reaction.

### NMR spectroscopy and data analysis

NMR experiments were recorded on an Agilent DD2 600 MHz four-channel spectrometer equipped with a triple-resonance cryogenically cooled probe-head or a Varian Unity Inova 500 MHz spectrometer equipped with a room-temperature triple-resonance probe. NMR samples contained 0.25 mM (for peptide K_D_ experiments) or 1 mM (all other experiments) CypA in 50 mM Na_2_HPO_4_, pH 6.5, 1 mM TCEP, 0.02% NaN3 and 10 (v/v) % D_2_O. The CypA S99T + AFPF sample contained ~0.85 mM CypA and 10.5 mM Suc-AFPF-pNA. Sample temperatures were calibrated using the 4% methanol + 96% methanol-d_4_ sample (DLM-5007, Cambridge Isotope Laboratories).

All data sets were processed with the NMRPipe/NMRDraw software package^40^ and visualized/analysed using the program NMRFAM-SPARKY^41^.

### Backbone assignments

TROSY-versions of a 3D HNCACB^42^ and CBCA(CO)NH^43^ experiments were recorded at 25 °C to obtain a nearly complete sequential backbone resonance assignment (H^N^, N, C^α^, C^β^) of CypA S99T/C115 and S99T/C115S/I97V. The HNCACB experiments were acquired with 50 (^15^N) × 70 (^13^C) × 537 (^1^H) complex points, with maximum evolution times equal to 22.2 (^15^N) × 8.3 (^13^C) × 64.0 (^1^H) ms. An interscan delay of 1 s was used with 8 scans per transient, giving rise to a net acquisition time of 36 h. The CBCA(CO)NH experiments were acquired with 59 (^15^N) × 50 (^13^C) × 537 (^1^H) complex points, with maximum evolution times equal to 26.2 (^15^N) × 5.9 (^13^C) × 64.0 (^1^H) ms. An interscan delay of 1 s was used with 8 scans per transient, giving rise to a net acquisition time of 30 h. Cross peak assignments were propagated to CPMG and CEST experiments at lower temperatures using [^1^H-^15^N]-TROSY-HSQC^44,45^ spectra recorded at different temperatures between 10 and 25 °C and where necessary confirmed using a 3D ^15^N-edited NOESY^46^ data set recorded at 10 °C. Cross peaks for Val2 and Glu81 were not visible at 25 °C, but could be assigned from the data recorded at 10 °C. The chemical shift assignments of CypA S99T/C115S and S99T/C115S/I97V have been deposited in the BioMagResBank^47^ with accession codes 27217 and 27218, respectively.

### K_D_ measurements

Dissociation constants for Suc-AFPF-pNA at 10 °C were obtained by titrating the peptide (final concentrations: 0, 1, 3, 5, 9 mM) into a solution of 0.25 mM CypA. Line-shape fitting was performed using PINT^48^ to obtained cross peak positions in the individual spectra. The combined chemical shift difference Δδ was calculated according to:

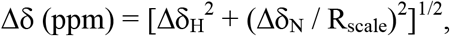

where *R*_scale_ = 6.3 was determined according to Mulder *et al.*^49^. Resonances with sufficient signal-to-noise and for which Δδ ≥ 0.035 ppm were included in the fits to determine dissociation constants using the following equation:

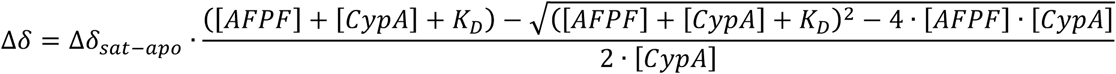

where [CypA] is the total enzyme concentration, [AFPF] is the concentration of peptide and Δδ_sat-apo_ is the combined chemical shift difference between apo and saturated CypA.

All resonances (13, 23, and 26 for CypA S99T, S99T/C115S and S99T/C115S/I97V, respectively) were fit simultaneously in Mathematica 11^50^ and standard errors are obtained from the global fit. The solubility of the peptide and sample stability limits the highest achievable concentration in our titration experiments and we could only attain data up to 5 mM AFPF for the double- and triple-mutant.

### CPMG relaxation dispersion experiments

Relaxation dispersion experiments on CypA were recorded on a 600 MHz spectrometer at different temperatures (10 and 15 °C for all mutants, and additionally at 20 °C for the triple-mutant). A TROSY-version of the relaxation-compensated ^15^N-CPMG pulse sequence^51,52^ was used, with the CPMG period implemented in a constant-time manner53.

The experiments were recorded as a series of 16 interleaved 2D data sets, with the constant-time relax period set to 40 ms. The CPMG field strengths were equal to 100, 200, 300, 400, 500, 600, 700, 800, 900, 950, and 1000 Hz, with duplicate experiments recorded for the reference experiment and 200, 800 and 1000 Hz. Experiments were acquired with 128 – 155 (^15^N) × 536 (^1^H) complex points, with maximum evolution times equal to 56.9 – 68.9 (^15^N) × 63.9 (^1^H) ms. An interscan delay of 2 s was used with 16, 24, or 32 scans per transient, giving rise to net acquisition times between 40 – 88 h for a complete pseudo-3D data set.

Line-shape fitting was performed using PINT^48^ and the obtained cross peak volumes were used to calculate the values of R_2,eff_. Error estimation in the experimental data using the four duplicate data points was performed as described earlier by Mulder *et al.*^54^. For ease of comparison, the values of R_2,inf_ were calculated by taking the average of the R_2,eff_ at the three highest v_CPMG_ values (i.e., 900, 950 and 1000 Hz) and normalized to the lowest temperature.

### CEST experiments

^15^N-CEST experiments^20^ on CypA were recorded on a 500 MHz spectrometer at 10 °C for two (single- and triple-mutants) or three (double-mutant) different ^15^N B_1_ field strengths. The weak irradiation fields applied during the relaxation delay were calibrated from 1D spectra as described by Guenneugues *et al.*^55^ with the irradiation position set to an isolated cross peak that did not show exchange.

The experiments on CypA S99T were recorded with a ^1^H decoupling field strength of 2.3 kHz (using 90_x_240_y_90_x_ composite pulses) during the relaxation delay, T_EX_, of 0.5 s. Two different ^15^N B_1_ fields, 16.3 and 29.7 Hz, were used and a series of 2D data sets were acquired with ^15^N offsets ranging between 96.5 (97.7) and 140.4 (139.2) ppm, in 112 (85) increments of 20 (25) Hz for n1 = 16.3 Hz (n1 = 29.7 Hz), and one reference experiment. Each 2D data set comprised of 110 (^15^N) × 512 (^1^H) complex points, with maximum evolution times equal to 52.4 (^15^N) × 64 (^1^H) ms, an interscan delay of 1.5 s and eight scans per transient were used, giving rise to net acquisition times of about 118 h (v_1_ = 16.3 Hz) and 94 h (v_1_ = 29.7 Hz).

For CypA S99T in the presence of Suc-AFPF-pNA, two different ^15^N B_1_ fields, 12.1 and 22.2 Hz, were used and a series of 2D data sets were acquired with ^15^N offsets ranging between 98.7 and 138.5 ppm, in 73 increments of 28 Hz, and one reference experiment. The experiments were recorded with a ^1^H decoupling field strength of 2.25 kHz (using 90_x_240_y_90_x_ composite pulses) during the relaxation delay, T_EX_, of 0.4 s (n1 = 22.2 Hz) or 0.5 s (v_1_ = 12.1 Hz). Each 2D data set comprised of 110 (^15^N) × 512 (^1^H) complex points, with maximum evolution times equal to 52.4 (^15^N) × 64 (^1^H) ms, an interscan delay of 1.5 s and twelve scans per transient were used, giving rise to net acquisition times of about 112 h (v_1_ = 22.2 Hz) and 116 h (v_1_ = 12.1 Hz).

For CypA S99T/C115S, three different ^15^N B_1_ fields, 12.2, 23.6, and 34.7 Hz, were used and a series of 2D data sets were acquired with ^15^N offsets ranging between 97.8 and 139.2 ppm, in 106 increments of 20 Hz, and one reference experiment. The experiments were recorded with a ^1^H decoupling field strength of 2.3–2.4 kHz (using 90_x_240_y_90_x_ composite pulses) during the relaxation delay, T_EX_, of 0.4 s (n1 = 23.6 Hz) or 0.5 s (n1 = 12.2 and 23.6 Hz). Each 2D data set comprised of 110 (^15^N) × 512 (^1^H) complex points, with maximum evolution times equal to 52.4 (^15^N) × 64 (^1^H) ms, an interscan delay of 1.5 s and eight scans per transient were used, giving rise to net acquisition times of about 107 h (v_1_ = 23.6 Hz) and 112 h (v_1_ = 12.2 and 23.6 Hz).

The experiments on CypA S99T/C115S/I97V were recorded with a ^1^H decoupling field strength of 2.4 kHz (using 90_x_240_y_90_x_ composite pulses) during the relaxation delay, T_EX_, of 0.4 s. Two different ^15^N B1 fields, 10.8 and 22.0 Hz, were used and a series of 2D data sets were acquired with ^15^N offsets ranging between 97.8 and 139.2 ppm, in 106 (85) increments of 20 (25) Hz for n1 = 10.8 Hz (n1 = 22.0 Hz), and one reference experiment. Each 2D data set comprised of 110 (^15^N) × 512 (^1^H) complex points, with maximum evolution times equal to 52.4 (^15^N) × 64 (^1^H) ms, an interscan delay of 1.5 s and eight scans per transient were used, giving rise to net acquisition times of about 86 h (v_1_ = 22.0 Hz) and 107 h (v_1_ = 10.8 Hz).

### CEST data analysis

Line-shape fitting was performed using PINT^48^ and the obtained cross peak volumes were used to calculate the ratio I / I_0_. The ^15^N-CEST profiles (ratio versus irradiation position) were analysed using the Python package ChemEx v0.6 (available from https://github.com/gbouvignies/chemex), which numerically propagates the Bloch-McConnell equations^56^, as described earlier^20^. Uncertainties in the ratio I / I_0_ were estimated from the apparent scatter in the baseline of the CEST profiles, whereas uncertainties in the fitting parameters (i.e., rate constants, populations and chemical shift differences) were determined from either the covariance matrix or 400-500 Monte-Carlo runs.

We analysed all the CEST profiles assuming the absence of exchange, a two-site exchange model and, if required, a three-site exchange model (only for CypA S99T/C115S and S99T/C115S/I97V). In the “no-exchange” situation, the values of k_ex_, p_B_, Δɷ and ΔR_2_ were fixed at 0 and only R_1_, R_2_ and I_0_ were fit on a per-residue basis. Residues that showed elevated R_2_ values and for which the χ^2^_red_ was significantly above 1, are likely candidates to experience an exchange process (Extended Data Figs. 3b, 4f and 5f).

For the two-site exchange model, initially only residues that showed a clear second dip or asymmetry were included. A per-residue fit was performed where in addition to the parameters above also k_ex_, p_B_ and Δɷ were allowed to float.

The clustering of k_ex_/p_B_ values for this initial subset indicated a single exchange process for CypA S99T (Extended Data Fig. 3a), and a global fit was performed after updating the initial k_ex_, p_B_ and residue-specific Δɷ values. As a third step, we fixed the global exchange parameters, (k_ex_ and p_B_) and re-fitted all residues. Residues that had a fitted value of |Δɷ| ≥ 1 ppm, an improved χ^2^_red_ and consistent fitting parameters in Monte-Carlo runs (not fixing k_ex_ and p_B_), were included in the list of probes experiencing exchange. Finally, a global fit and 500 Monte-Carlo runs were performed with every exchanging residue and now allowing all parameters [k_ex_/p_B_ (global) and Δɷ, R_1_, R_2_, DR_2_ and I_0_ (residue-specific)] to float. The results of this Monte-Carlo analysis are shown in Extended Data Fig. 3c for k_ex_/p_B_ and chemical shift differences are plotted on the structure (Extended Data Fig. 3d). Furthermore, assuming a two-state exchange model, the χ^2^_red_ and fitted R_2_ values for exchanging residues went down to their expected values, with the exception of the loop region undergoing exchange on the millisecond timescale that still exhibits high R_2_ values (Extended Data Fig. 3b).

On the contrary, for CypA S99T/C115S and S99T/C115S/I97V the clustering of k_ex_/p_B_ values for the initial subset suggested the presence of two different processes (Extended Data Figs. 4a and 5a). Residues that seemingly fitted well to a two-site exchange model were grouped into two clusters and separately fitted to a global process, yielding two different rates of interconversion and populations (Extended Data Figs. 4b and 5b). In a similar manner as for CypA S99T we separately fixed the two combinations of global exchange parameters and re-fitted all data. Residues that had a fitted value of |Δɷ| ≥ 1 ppm, an improved χ^2^_red_ and consistent fitting parameters in Monte-Carlo runs (not fixing k_ex_ and p_B_), were included in the appropriate cluster of probes experiencing exchange. The two clusters were obtained consistently even when k_ex_/p_B_ values from the other one were used as starting points of the minimization. The residues present in the clusters are found in two different parts of the protein (*cf.* green and red spheres in Fig. 2e,g) and also have different Rex contributions in the CPMG relaxation dispersion profiles (Supplementary Data 1 and 2). Furthermore, CEST data for several residues, including F88 and N102, could not be fit to a two-site exchange model (Extended Data Figs. 4c and 5c).

Taken together, these observations show that the CEST data for the double- and triple-mutant require a three-state exchange model to explain the data. We note that for some residues (*cf.* orange spheres in Fig. 2e,g), the fast loop motion and slow processes give rise to convoluted exchange data. However, this does not automatically mean that any (or all) of these processes are correlated; it can simply be that a probe participating in one exchange process “feels” the fluctuating field of another, nearby exchange phenomena.

There are several possibilities to connect three different states (A=ground state, B=minor state 1, and C=minor state 2): one triangular model, where all states are connected, and three linear models (A⇔B⇔C, A⇔C⇔B, and B⇔A⇔C) as described earlier by Sekhar *et al.*^57^. We did not consider the triangular model as it requires one additional fitting parameter and our NMR data is not of high enough quality to warrant this. From fitting the individual residues, we already obtained two values for k_ex_/population (Extended Data Figs. 4b and 5b) and Δω values, and those were used as starting values in the linear models. The combination with highest population was used as k_ex,AB_/p_B_/Δω_AB_ and the other as k_ex,AC_/p_C_/Δω_AC_. The ACB-model, where the least populated minor state is the intermediate, did not fit the data; whereas the other two models gave very similar results and seem to describe the data equally well. Based on our current NMR and crystallography experiments and earlier data on CypA S99T^11^, we favor the B⇔A⇔C model, where both processes are independent and this model has been used for further analysis.

For a number of CEST profiles, including F88 and N102, we do see two dips/asymmetry in the CEST traces (Extended Data Figs. 4d and 5d) and can, therefore, determine the chemical shift of all three states. The CEST data for these residues could be fitted to the BAC-model, with rate constants and populations very similar obtained for the “pure” two-state exchange models, indicating that the observed profiles are caused by these two, independent, slow-exchange processes. For others only one dip was visible, and there we determined the best initial value for Dwof the second minor state by starting from the two possible options (i.e., the chemical shift of the other minor state is either close to the major state or to observable minor state) and comparing the χ^2^_red_ value.

Finally, a global fit and 400-500 Monte-Carlo runs were performed with every exchanging residue and now allowing all parameters [kex,AB/p_B_; k_ex,AC_/p_C_ (global) and Δω_AB_,Δω_AC_, R_1_, R_2_, ΔR_2_ and I_0_ (residue-specific)] to float. The results of this Monte-Carlo analysis are shown in Extended Data Figs. 4e and 5e for k_ex_/population and chemical shift differences are plotted on the CypA structure (Extended Data Figs. 4f and 5f). Furthermore, assuming a three-state exchange model, the χ^2^_red_ and fitted R_2_ values for exchanging residues went down to their expected values, with the exception of the loop region undergoing exchange on the millisecond timescale that still exhibits high R_2_ values (Extended Data Figs. 4f and 5f).

## Data Availability

Structure factors and refined model of CypA S99T/C115S/I97V has been deposited in the PDB under accession code 5WC7. The NMR assignments of CypA S99T/C115S and S99T/C115S/I97V have been deposited in the BMRB under accession codes 27217 and 27218, respectively.

## Acknowledgements

We thank N. Ollikainen, T. Kortemme and H. van den Bedem for discussions. We are grateful to L. Kay, R. Muhandiram and G. Bouvignies for sharing their NMR pulse sequence codes and help with the ChemEx software. F. Pontiggia is acknowledged for his help to implement the NMR data fitting workflow on the Brandeis HPC cluster. We thank G. Coaker for AvrRpt2 constructs. This work was supported by the Howard Hughes Medical Institute, the Office of Basic Energy Sciences, Catalysis Science Program, U.S. Dept. of Energy, award DE-FG02-05ER15699 and NIH (GM100966) to D.K, and a Searle Scholar Award from the Kinship Foundation, a Pew Scholar Award from the Pew Charitable Trusts, a Packard Fellowship from the David and Lucile Packard Foundation, NIH (GM110580), and NSF (STC-1231306) to J.S.F. R.O. was supported as an HHMI Fellow of the Damon Runyon Cancer Research Foundation (DRG-2114-12). Data collection at BL831 at the Advanced Light Sources is supported by the Director, Office of Science, Office of Basic Energy Sciences, of the U.S. Department of Energy under Contract No. DE-AC02-05CH11231, UC Office of the President, Multicampus Research Programs and Initiatives grant MR-15-32859, and the Program Breakthrough Biomedical Research, which is partially funded by the Sandler Foundation.

## Author Contributions

R.O., L.L., D.K. and J.S.F. designed experiments. L.L. and D.M. performed the library screen and developed the assay for screening activity in cell lysate with supervision of D.S.T. and J.S.F. L.L. performed the activity assay and R.O. analysed the data. L.L., L.R.K. and J.S.F. performed the X-ray experiments; R.O. and M.W.C. performed the NMR experiments and R.O. analysed the data. R.O., J.S.F. and D.K. wrote the paper. All authors contributed to data interpretation and commented on the manuscript. Correspondence and requests for materials should be addressed to J.S.F. or D.K.

---

**Extended Data Figure 1.**
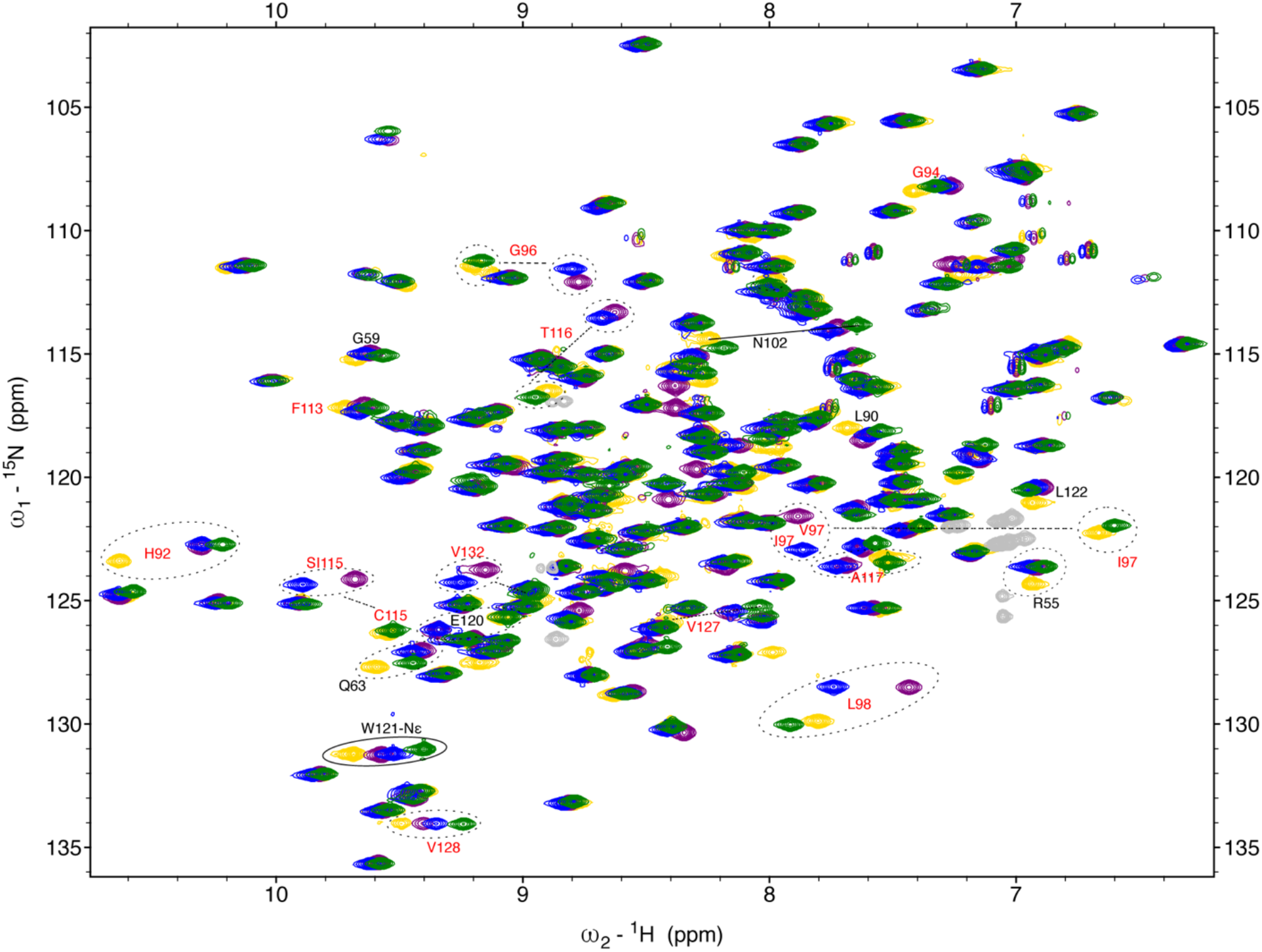
Overlay of ^15^N-[TROSY]-HSQC spectra for wild-type CypA (yellow), S99T (green) and rescue mutants (S99T/C115S, blue; S99T/C115S/I97V, purple). All spectra were recorded on ~1 mM protein samples using a 600 MHz spectrometer at 10 °C. Sequence-specific assignments given in red indicate residues that have moved significantly due to proximity, in sequence and/or space, to the mutation site (<5 Å). Cross peaks connected with a solid line or in a solid circle show the population inversion upon the Ser99 mutation and the partial shift towards wild-type for the rescue mutants. Aliased Arg-Nε side chain signals are shown in grey for all spectra.

**Extended Data Figure 2.**
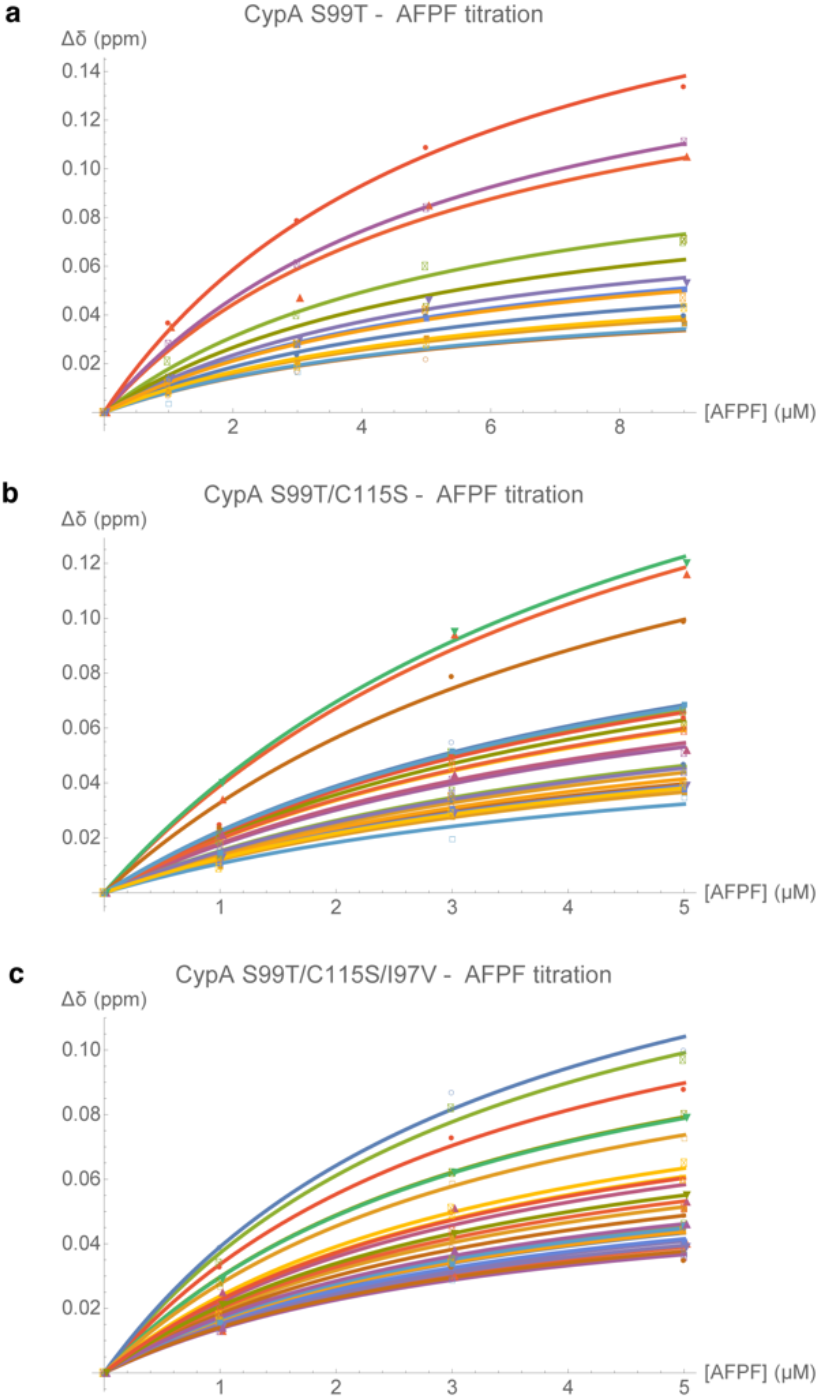
K_D_ determination for the three mutant forms of CypA for Suc-AFPF-pNA measured by NMR chemical shift analysis from peptide titrations. Resonances for which Δδ ≥ 0.035 ppm (thirteen for CypA S99T **(a)**, twenty-three for S99T/C115S **(b)** and twenty-six for S99T/C115S/I97V **(c)**, respectively) were fit simultaneously in Mathematica 11^50^ and standard errors are obtained from the global fit.

**Extended Data Figure 3.**
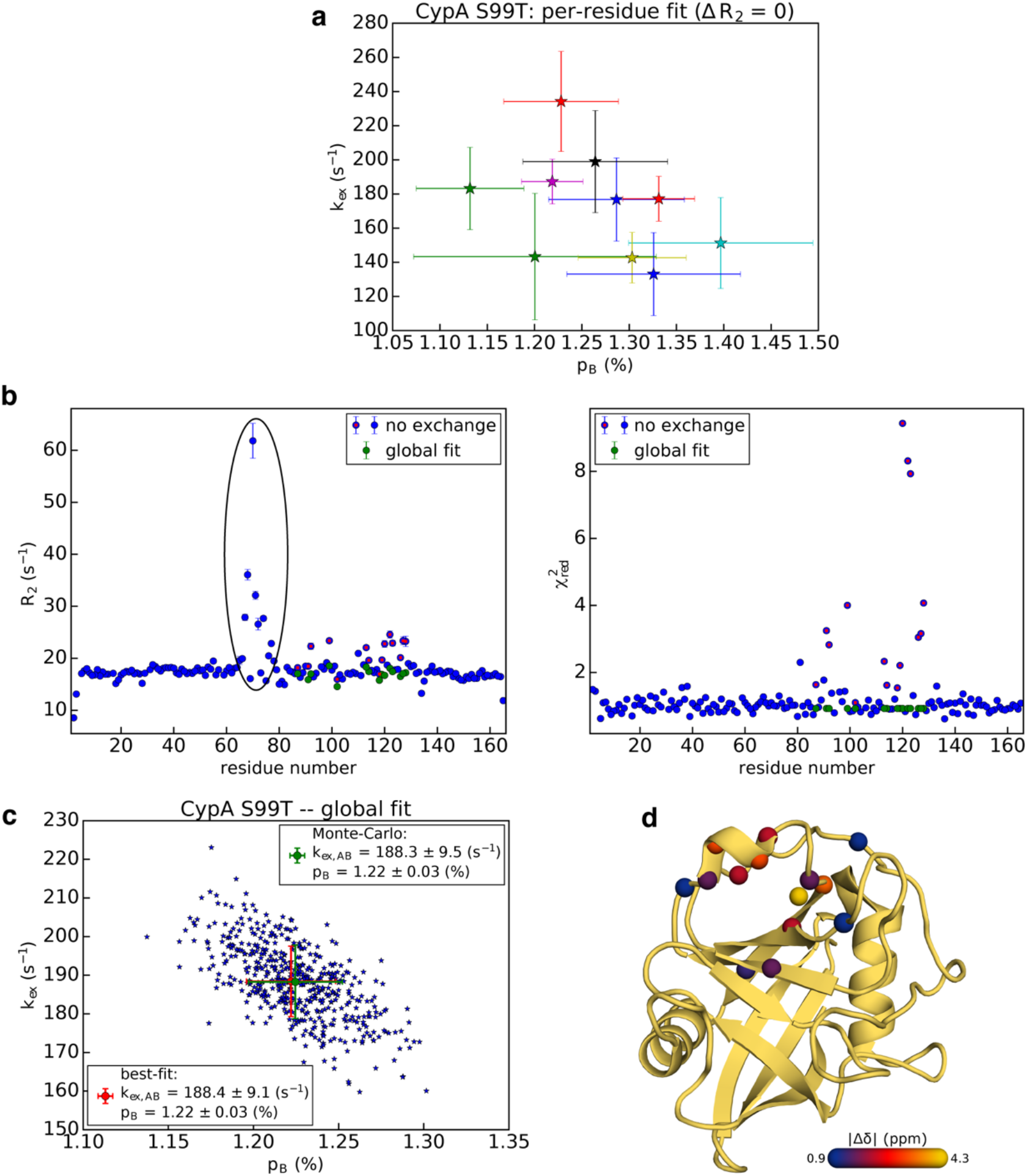
Analysis of the ^15^N-CEST data for CypA S99T. (a) Per-residue fit of the CEST profiles for the initial 10 residues that clearly show slow exchange by visual inspection. Clustering of the k_ex_/p_B_ values indicate the presence of one global exchange process. **(b)** Comparison of R_2_ and χ^2^_red_ values for no-exchange model (blue) and one global, slow-exchange process (green). For ease of comparison residues that are included in the global fit are also shown with a red marker for the no-exchange model. Fitting the data assuming no exchange results in elevated R_2_ values (left) for the loop region (residues 65-80), whereas χ^2^_red_ values (right) are close to one. This observation is consistent with line broadening of the ground-state signal due to dynamics on the millisecond timescale and the absence of a second dip or asymmetry that would indicate a slow-exchange process. In contrast, assuming a no-exchange model for residues that, in fact, do experience a slow-exchange process, results in large χ^2^_red_ values in addition to a higher R_2_ in an attempt to –unsuccessfully– explain the asymmetry in the CEST profiles. The data show that the data can be explained satisfactory by one, global exchange process as judged from the R_2_ and χ^2^_red_ values. **(c)** Uncertainties in the fitting parameters for the global fit were determined from 500 Monte-Carlo simulations and the resulting scatter plot for k_ex_ vs. p_B_ is shown. The values and their uncertainties obtained using this method and estimated from the covariance matrix are nearly identical. **(d)** The chemical shift difference between the states, |Dd|, for the 15 residues is plotted on the structure of CypA.

**Extended Data Figure 4.**
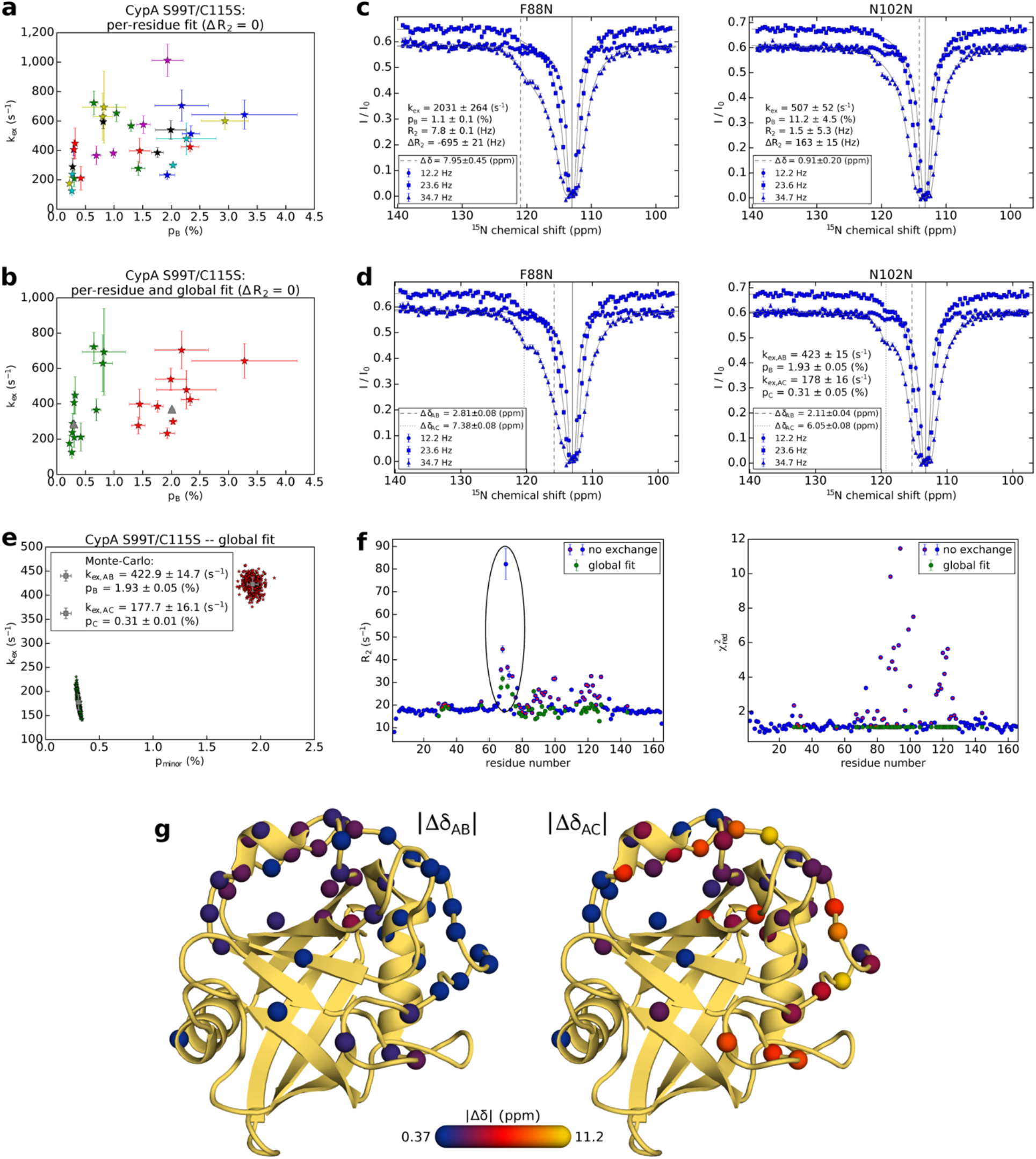
Analysis of the ^15^N-CEST data for CypA S99T/S115S indicates the presence of more than one exchange process. (a) Per-residue fit of the CEST profiles for the initial 30 residues that clearly show slow exchange by visual inspection. Clustering of the k_ex_/p_B_ values suggests the presence of two distinct exchange processes. **(b)** Residues that are well-described by a two-state exchange model were split into two clusters (*c.f.*, red (11 residues) and green (22 residues) spheres in Fig. 2e). Here, only the residues that are present in panel a and are used in the two-state global fit are shown together with their respective global k_ex_/p_B_ values (grey triangles). (c-d) Several residues, including F88 and N102, do not fit to a two-site exchange process even if all fitting parameters are allowed to float **(c)**. However, a three-site exchange model can explain the experimental data **(d)** and the vales of k_ex_/p_minor_ correspond well to the two observed clusters assuming a single two-site exchange model (panel b). **(e)** Uncertainties in the fitting parameters for the three-state global fit that include all 46 residues were determined from 400 Monte-Carlo simulations and the resulting scatter plot for k_ex_ vs. p_minor_ is shown. The values and their uncertainties obtained using this method and from the covariance matrix are nearly identical. **(f)** Residue-specific values of R_2_ and χ^2^_red_ assuming there is no exchange (blue) or a global, three-site process (green). For ease of comparison, the 46 residues that experience exchange and are included in the global fit are also shown with a red marker for the no-exchange model. Fitting a no-exchange model shows significantly elevated R_2_ and χ^2^_red_ values for a large number of residues. After fitting an appropriate exchange model R_2_ and χ^2^_red_ drop to expected values, with the exception of the loop region (residues 65-80), where elevated R_2_ values are still observed, consistent with line broadening of the ground-state signal due to dynamics on the millisecond timescale. **(g)** The chemical shift differences between the states, |Δδ_AB_| (left panel) and |Δδ_AC_| (right panel), for the 46 residues are plotted on the structure of CypA.

**Extended Data Figure 5.**
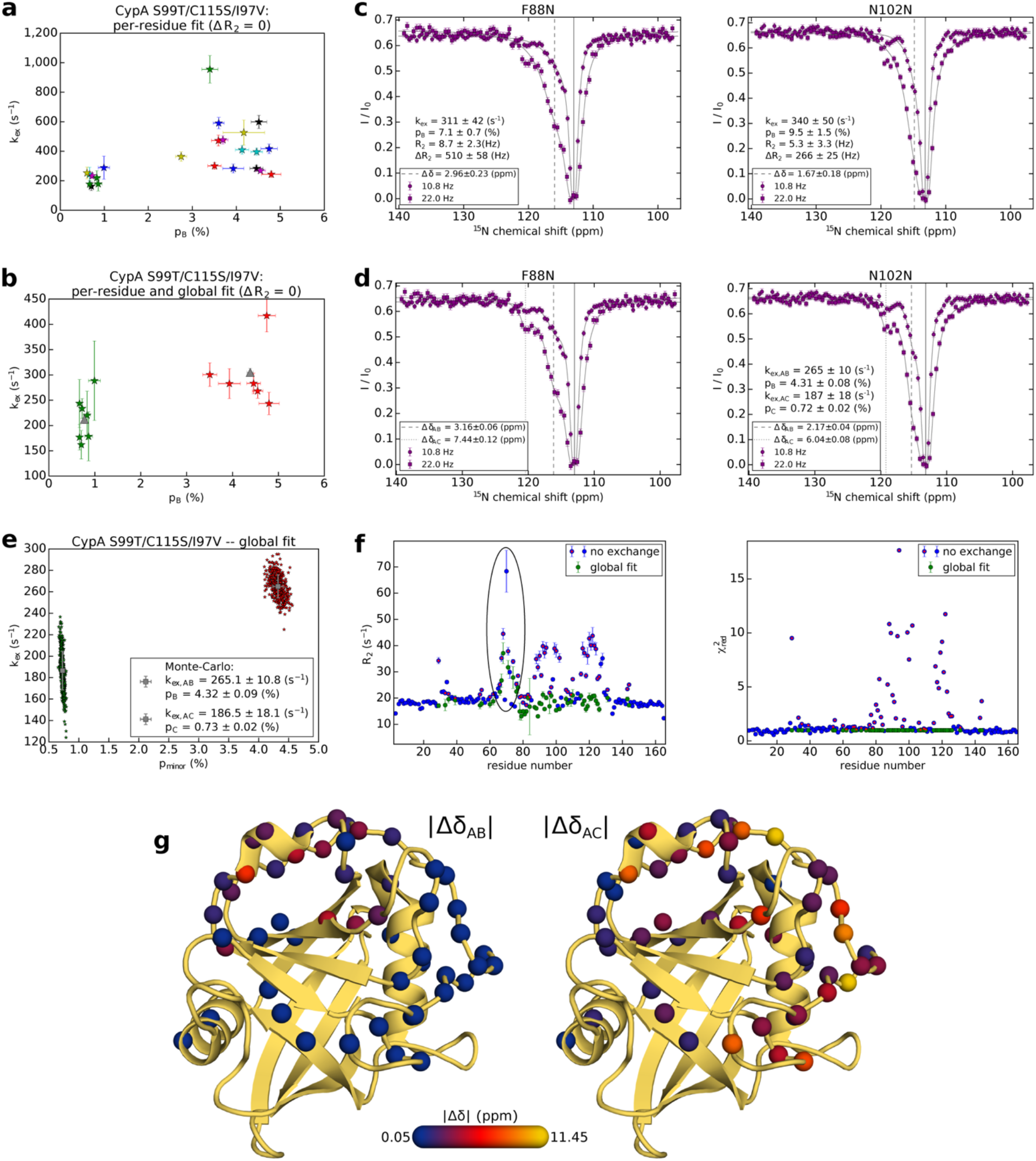
Analysis of the ^15^N-CEST data for CypA S99T/S115S/I97V indicates the presence of more than one exchange process. (a) Per-residue fit of the CEST profiles for the initial 23 residues that clearly show slow exchange by visual inspection. Clustering of the k_ex_/p_B_ values suggest the presence of two distinct exchange processes. **(b)** Residues that are well-described by a two-state exchange model were split into two clusters (*c.f.*, red (12 residues) and green (25 residues) spheres in Fig. 2g). Here, only the residues that are in panel a and are used in the two-state global fit are shown together with their respective global k_ex_/p_B_ values (grey triangles). (c-d) Several residues, including F88 and N102, do not fit to a two-site exchange process even if all fitting parameters are allowed to float **(c)**. However, a three-site exchange model can explain the experimental data **(d)** and the vales of k_ex_/p_minor_ correspond well to the two observed clusters assuming a single two-site exchange model (panel b). **(e)** Uncertainties in the fitting parameters for the three-state global fit that include all 55 residues were determined from 425 Monte-Carlo simulations and the resulting scatter plot for k_ex_ vs. p_minor_ is shown. The values and their uncertainties obtained using this method and from the covariance matrix are nearly identical. **(f)** Residue-specific values of R_2_ and χ^2^_red_ assuming there is no exchange (blue) or a global, three-site process (green). For ease of comparison, the 55 residues that experience exchange and are included in the global fit are also shown with a red marker for the no-exchange model. Fitting a no-exchange model shows significantly elevated R_2_ and χ^2^_red_ values for a large number of residues. After fitting an appropriate exchange model R_2_ and χ^2^_red_ drop to expected values, with the exception of the loop region (residues 65-80), where elevated R_2_ values are still observed, consistent with line broadening of the ground-state signal due to dynamics on the millisecond timescale. **(g)** The chemical shift differences between the states, |Δδ_AB_| (left panel) and |Δδ_AC_| (right panel), for the 55 residues are plotted on the structure of CypA.

**Extended Data Figure 6.**
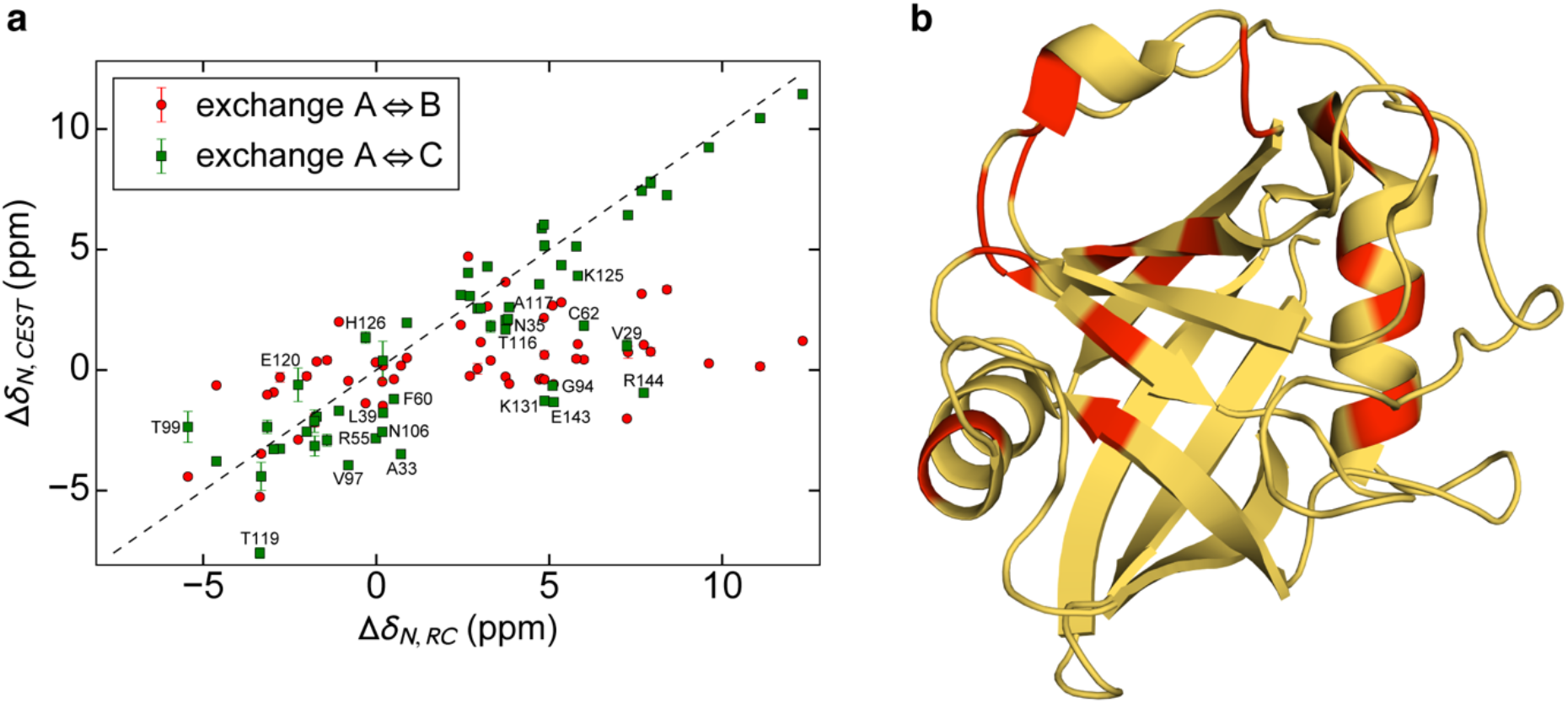
Correlation between ^15^N Δδ values obtained from CEST and those between the major state and predicted random coil chemical shifts of CypA S99T/C115S/I97V. **(a)** ^15^N chemical shift differences between state A⇔B (red circles) and A⇔C (green squares) were extracted from the 3-state global fit of the CEST profiles, and the random coil chemical shifts were predicted using the method of Tamiola *et al.*^58^. There is no clear correlation between Δδ_N,AB_ and Δδ_N,RC_ (pairwise rmsd = 4.2 ppm). On the contrary, the correlation between Δδ_AC_ and Δδ_RC_ suggests that minor state C corresponds to a more extended/unfolded conformation. Residues that are labeled with their assignment are not correlated (|Δδ_N,AC_-Δδ_N,RC_| ≥ 1.5 ppm), indicating that these do not sample an extended conformation, and are color in red on the structure **(b)**. The majority of these residues belong to group-I or are part of the dynamic network that is involved in the catalysis. The pairwise rmsd for Δδ_N,AC_ and Δδ_N,RC_ is 2.6 ppm when considering all residues and 0.9 ppm excluding the labeled ones that are part of the dynamic network.

**Extended Data Figure 7.**
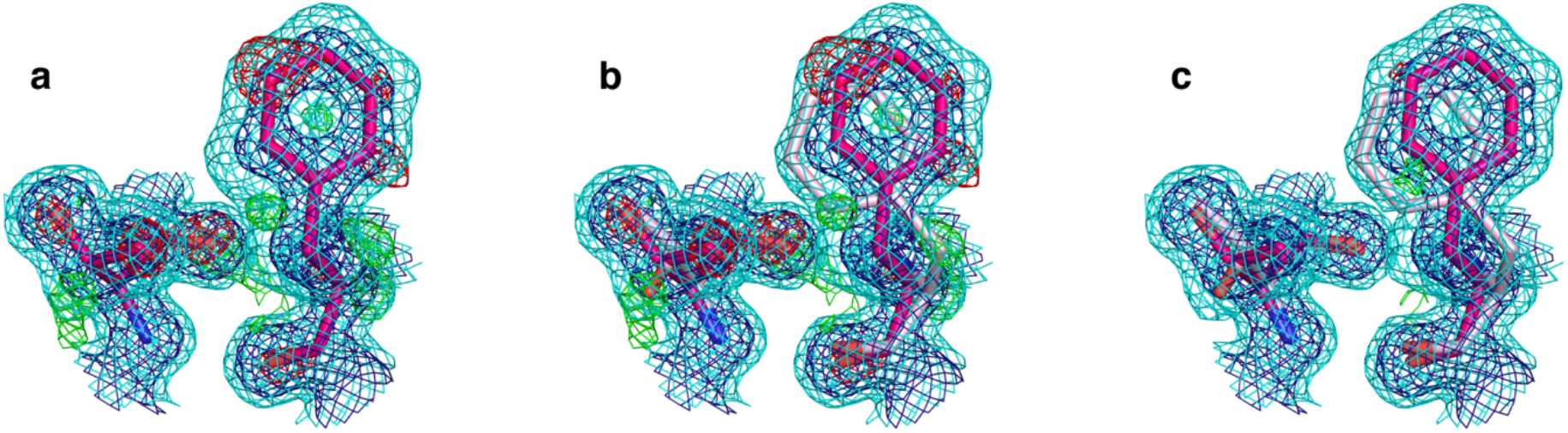
Difference electron density for alternative conformations of Thr99 and Phe113 in CypA S99T/C115S/I97V. **(a)** Negative and positive difference peaks in mFo-DFc electron density (±3.0σ in green and red) are contained within expanded 2mFo-DFc electron density (2.0σ in dark blue and 0.3σ in cyan), and not accounted for by a single conformer model (magenta) of CypA S99T/C115S/I97V. **(b)** The electron density maps as shown in **(a)** with the refined qFit multiconformer model (alternative conformations in magenta, ~0.8 occupancy, and light pink, ~0.2 occupancy) that explains the difference features. **(c)** The refined qFit models shown in **(b)** with the final 2mFo-DFc (2.0σ in dark blue and 0.3σ in cyan) and mFo-DFc maps (±3.0σ in green and red).

**Extended Data Figure 8.**
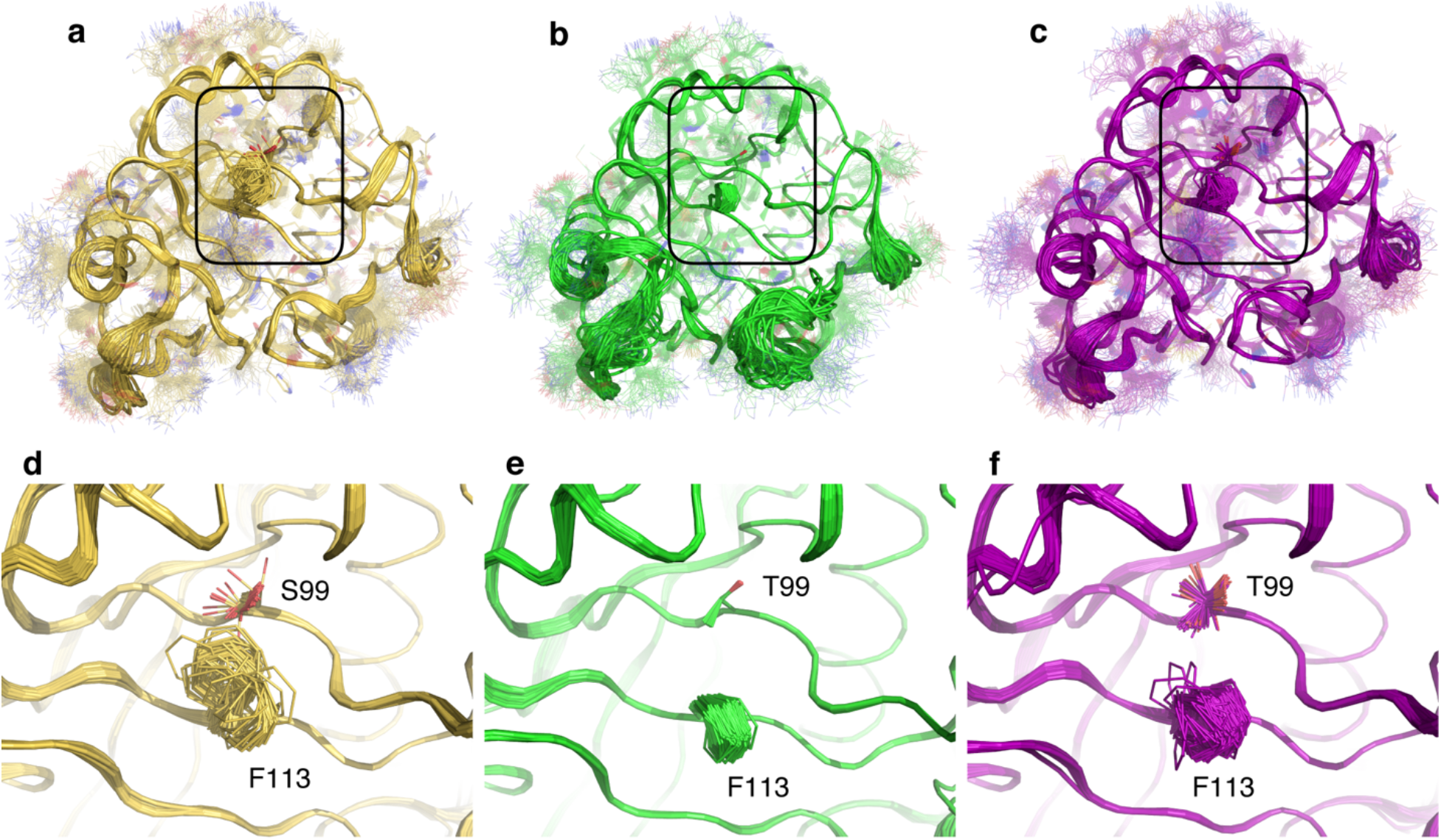
Time-averaged ensemble refinement for wild-type CypA (yellow), S99T (green), and S99T/C115S/I97V (purple). All refinements were performed using phenix.ensemble refinement^38^ with parameters (pTLS, wxray, tx) selected based on lowest Rfree. Overall view (a-c) and corresponding zoom-in **(d-f)** for wild-type CypA (a,d) (refined with pTLS 0.775, wxray 8.125, and tx 2.0) shows extensive side chain heterogeneity in the active site extending to the core through residues Phe113 and Ser99 (shown in sticks); S99T **(b,e)** (refined with pTLS 0.775, wxray 8.125, and tx 2.0) shows no heterogeneity in Phe113 and Thr99; and S99T/C115S/I97V **(c,f)** (refined with pTLS 0.55, wxray 8.125, and tx 2.0) shows increased heterogeneity in Phe113 and multiple conformations of Thr99 relative to the S99T.

**Extended Data Figure 9.**
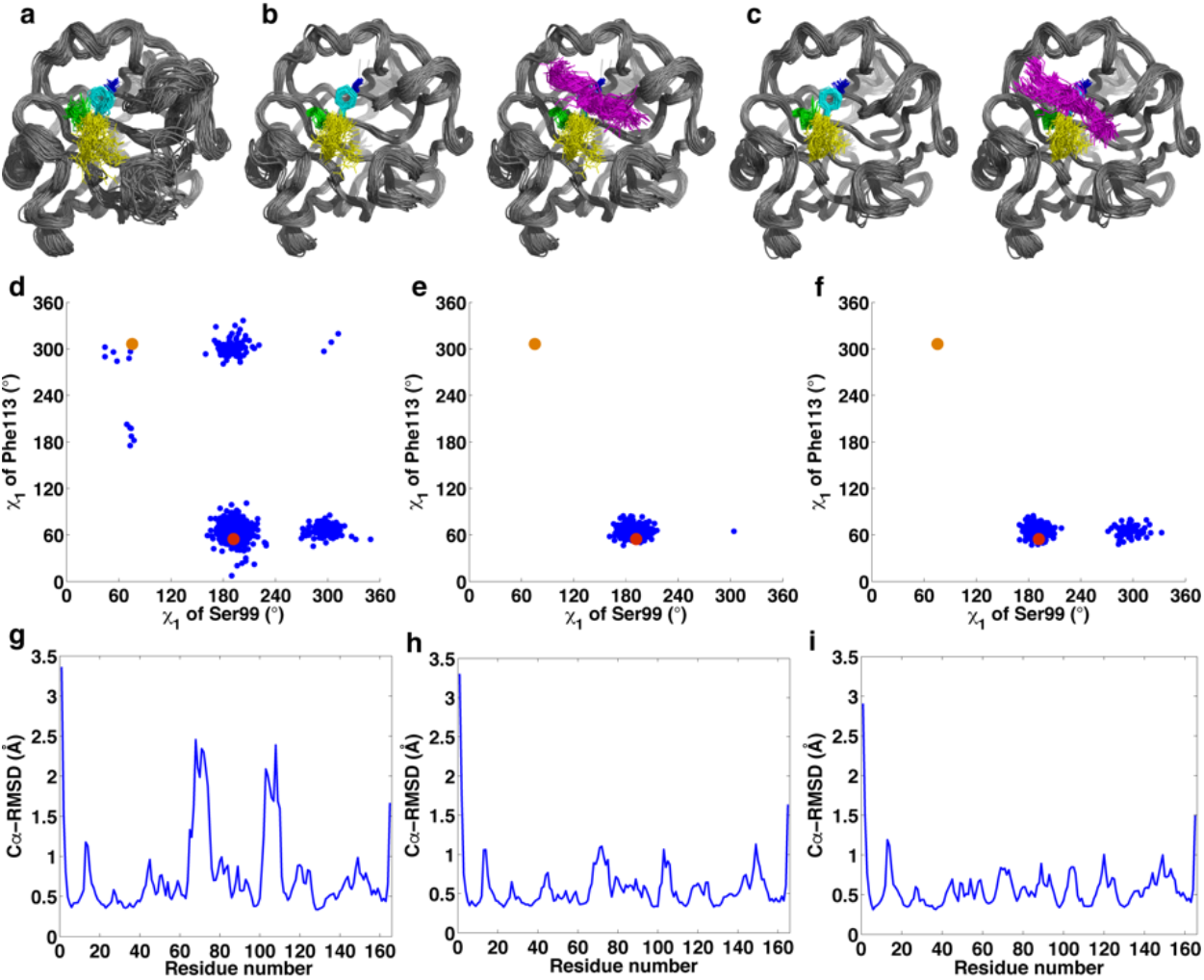
Analysis of conformational ensembles calculated by Camilloni *et al.* using DFT calculations to propose a catalytic mechanism^26^. Conformational heterogeneity is suppressed in the catalytic complexes, and alternate conformations seen in the X-ray structures^11^ are not sampled in the simulation with the substrates bound. **(a-c)** A subset of 68 randomly selected ensemble members is shown in cartoon with residues Ser99 (blue), Phe113 (cyan), Met61 (green), and Arg55 (yellow) shown in sticks. The apo protein ensemble **(a)** shows more conformational heterogeneity than the peptide (GSFGPDLRAGD) substrate-bound forms with a Gly-Pro in cis- **(b)** and trans- **(c)** conformations, shown without (left) or with (right) the substrate backbone (magenta). Curiously, their cis- and trans-peptide complexes show only minimal protein conformational differences compared to the starting crystal structures, while the free CypA showed multiple substates in their simulations. This is reflected in the dihedral angle distribution for Ser99 and Phe113 for apo wild-type CypA **(d)** versus the cis- **(e)** and trans- **(f)** substrate bound ensembles. The two alternative (major and minor) conformations seen in the room-temperature X-ray structure of wild-type CypA (PDB 3k0n) are shown in as red and orange dots for reference. Their result is in sharp contrast with our NMR dynamics for WT^10^ and S99T during catalysis measured here, which clearly shows that conformational substates interconvert across the core catalytic network and that this rate is correlated to catalysis. Although no significant change in loop dynamics is detected by NMR during catalysis, the MD simulations indicate larger deviations in the C^a^RMSDs in the apo protein **(g)** is specifically decreased in the loops surrounding residues 60-80 and 100-120 in the cis- **(h)** and trans- **(i)** substrate bound ensembles. New simulations that incorporate the side-chain dynamics characterized here may help bridging their DFT computational approach which emphasize a key role for electrostatics, with our experimental results that deliver the protein conformational substates.

**Extended Data Table 1.**
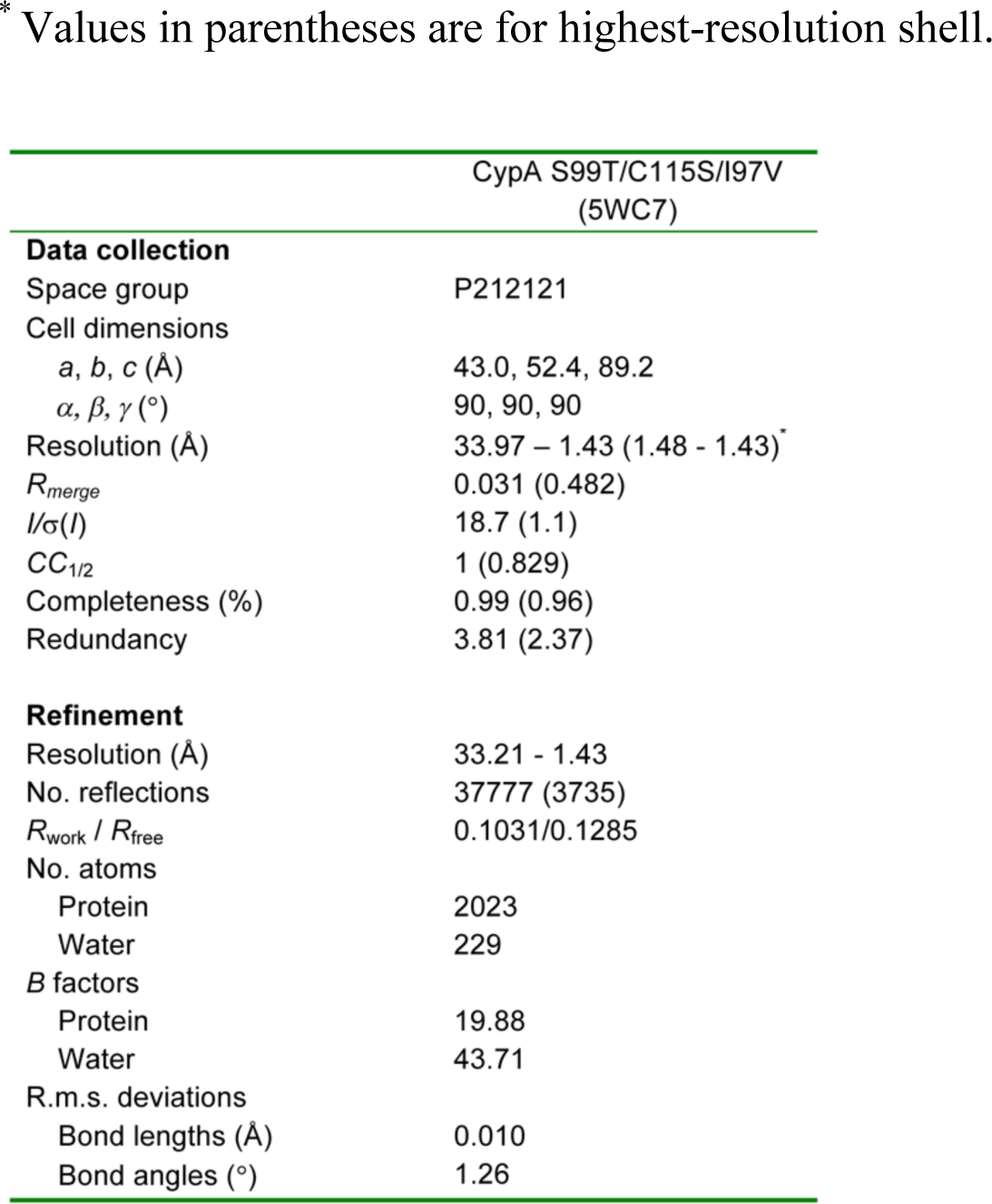
Data collection and refinement statistics. The number of crystals for each structure is 1.

## References

1 Blomberg, R. et al. Precision is essential for efficient catalysis in an evolved Kemp eliminase. Nature 503, 418–421, doi:10.1038/nature12623 (2013).

2 Romero, P. A. & Arnold, F. H. Exploring protein fitness landscapes by directed evolution. Nat Rev Mol Cell Biol 10, 866–876, doi:10.1038/nrm2805 (2009).

3 Arnold, F. H. The nature of chemical innovation: new enzymes by evolution. Q Rev Biophys 48, 404–410, doi:10.1017/S003358351500013X (2015).

4 Kay, L. E. New Views of Functionally Dynamic Proteins by Solution NMR Spectroscopy. J Mol Biol 428, 323–331, doi:10.1016/j.jmb.2015.11.028 (2016).

5 Boehr, D. D., Dyson, H. J. & Wright, P. E. An NMR perspective on enzyme dynamics. Chem Rev 106, 3055–3079, doi:10.1021/cr050312q (2006).

6 Smock, R. G. & Gierasch, L. M. Sending signals dynamically. Science 324, 198–203, doi:10.1126/science.1169377 (2009).

7 Palmer, A. G., 3rd. Enzyme dynamics from NMR spectroscopy. Acc Chem Res 48, 45–465, doi:10.1021/ar500340a (2015).

8 Shukla, D., Hernandez, C. X., Weber, J. K. & Pande, V. S. Markov state models provi insights into dynamic modulation of protein function. Acc Chem Res 48, 414–422, doi:10.1021/ar5002999 (2015).

9 Eisenmesser, E. Z., Bosco, D. A., Akke, M. & Kern, D. Enzyme dynamics during catalysis. Science 295, 1520–1523, doi:10.1126/science.1066176 (2002).

10 Eisenmesser, E. Z. et al. Intrinsic dynamics of an enzyme underlies catalysis. Nature 438, 117–121, doi:10.1038/nature04105 (2005).

11 Fraser, J. S. et al. Hidden alternative structures of proline isomerase essential for catalysis. Nature 462, 669–673, doi:10.1038/nature08615 (2009).

12 Harms, M. J. & Thornton, J. W. Evolutionary biochemistry: revealing the historical and physical causes of protein properties. Nat Rev Genet 14, 559–571, doi:10.1038/nrg35 (2013).

13 Ma, B. & Nussinov, R. Protein dynamics: Conformational footprints. Nat Chem Biol 12 890–891, doi:10.1038/nchembio.2212 (2016).

14 Kern, D., Kern, G., Scherer, G., Fischer, G. & Drakenberg, T. Kinetic analysis of cyclophilin-catalyzed prolyl cis/trans isomerization by dynamic NMR spectroscopy. Biochemistry 34, 13594–13602 (1995).

15 Harrison, R. K. & Stein, R. L. Mechanistic studies of peptidyl prolyl cis-trans isomerase: evidence for catalysis by distortion. Biochemistry 29, 1684–1689 (1990).

16 Coaker, G., Zhu, G., Ding, Z., Van Doren, S. R. & Staskawicz, B. Eukaryotic cyclophilin as a molecular switch for effector activation. Mol Microbiol 61, 1485–1496, doi:10.1111/j.1365-2958.2006.05335.x (2006).

17 Aumuller, T., Jahreis, G., Fischer, G. & Schiene-Fischer, C. Role of prolyl cis/trans isomers in cyclophilin-assisted Pseudomonas syringae AvrRpt2 protease activation. Biochemistry 49, 1042–1052, doi:10.1021/bi901813e (2010).

18 Kofron, J. L., Kuzmic, P., Kishore, V., Colon-Bonilla, E. & Rich, D. H. Determination of kinetic constants for peptidyl prolyl cis-trans isomerases by an improved spectrophotometric assay. Biochemistry 30, 6127–6134 (1991).

19 Chi, C. N. et al. A Structural Ensemble for the Enzyme Cyclophilin Reveals an Orchestrated Mode of Action at Atomic Resolution. Angew Chem Int Ed Engl 54, 1165711661, doi:10.1002/anie.201503698 (2015).

20 Vallurupalli, P., Bouvignies, G. & Kay, L. E. Studying “invisible” excited protein states in slow exchange with a major state conformation. J Am Chem Soc 134, 8148–8161, doi:10.1021/ja3001419 (2012).

21 Keedy, D. A., Fraser, J. S. & van den Bedem, H. Exposing Hidden Alternative Backbone Conformations in X-ray Crystallography Using qFit. PLoS Comput Biol 11, e1004507, doi:10.1371/journal.pcbi.1004507 (2015).

22 van den Bedem, H., Bhabha, G., Yang, K., Wright, P. E. & Fraser, J. S. Automated identification of functional dynamic contact networks from X-ray crystallography. Nat Methods 10, 896–902, doi:10.1038/nmeth.2592 (2013).

23 Tokuriki, N. et al. Diminishing returns and tradeoffs constrain the laboratory optimization of an enzyme. Nat Commun 3, 1257, doi:10.1038/ncomms2246 (2012).

24 Nagel, Z. D. & Klinman, J. P. A 21st century revisionist’s view at a turning point in enzymology. Nat Chem Biol 5, 543–550, doi:10.1038/nchembio.204 (2009).

25 Kamerlin, S. C. & Warshel, A. At the dawn of the 21st century: Is dynamics the missing link for understanding enzyme catalysis? Proteins 78, 1339–1375, doi:10.1002/prot.22654 (2010).

26 Camilloni, C. et al. Cyclophilin A catalyzes proline isomerization by an electrostatic handle mechanism. Proc Natl Acad Sci U S A 111, 10203–10208, doi:10.1073/pnas.1404220111 (2014).

27 Papaleo, E., Sutto, L., Gervasio, F. L. & Lindorff-Larsen, K. Conformational Changes and Free Energies in a Proline Isomerase. J Chem Theory Comput 10, 4169–4174, doi:10.1021/ct500536r (2014).

28 Dodani, S. C. et al. Discovery of a regioselectivity switch in nitrating P450s guided by molecular dynamics simulations and Markov models. Nat Chem 8, 419–425, doi:10.1038/nchem.2474 (2016).

29 Zhao, Y. & Ke, H. Crystal structure implies that cyclophilin predominantly catalyzes the trans to cis isomerization. Biochemistry 35, 7356–7361, doi:10.1021/bi9602775 (1996).

30 Ke, H. Similarities and differences between human cyclophilin A and other β-barrel structures. Journal of Molecular Biology 228, 539–550, doi:10.1016/0022-2836(92)90841-7 (1992).

## References

31 Rockah-Shmuel, L., Toth-Petroczy, A. & Tawfik, D. S. Systematic Mapping of Protein Mutational Space by Prolonged Drift Reveals the Deleterious Effects of Seemingly Neutral Mutations. PLoS Comput Biol 11, e1004421, doi:10.1371/journal.pcbi.1004421 (2015).

32 Kabsch, W. Xds. Acta Crystallogr D Biol Crystallogr 66, 125–132, doi:10.1107/S0907444909047337 (2010).

33 Keedy, D. A. et al. Mapping the conformational landscape of a dynamic enzyme by multitemperature and XFEL crystallography. Elife 4, doi:10.7554/eLife.07574 (2015).

34 Karplus, P. A. & Diederichs, K. Linking crystallographic model and data quality. Science 336, 1030–1033, doi:10.1126/science.1218231 (2012).

35 McCoy, A. J. et al. Phaser crystallographic software. J Appl Crystallogr 40, 658–674, doi:10.1107/S0021889807021206 (2007).

36 Emsley, P., Lohkamp, B., Scott, W. G. & Cowtan, K. Features and development of Coot. Acta Crystallogr D Biol Crystallogr 66, 486–501, doi:10.1107/S0907444910007493 (2010).

37 Afonine, P. V. et al. Towards automated crystallographic structure refinement with phenix.refine. Acta Crystallogr D Biol Crystallogr 68, 352–367, doi:10.1107/S0907444912001308 (2012).

38 Burnley, B. T., Afonine, P. V., Adams, P. D. & Gros, P. Modelling dynamics in protein crystal structures by ensemble refinement. Elife 1, e00311, doi:10.7554/eLife.00311 (2012).

39 The PyMOL Molecular Graphics System v. 1.8.6 (2017).

40 Delaglio, F. et al. NMRPipe: a multidimensional spectral processing system based on UNIX pipes. J Biomol Nmr 6, 277–293, doi:Doi 10.1007/Bf00197809 (1995).

41 Lee, W., Tonelli, M. & Markley, J. L. NMRFAM-SPARKY: enhanced software for biomolecular NMR spectroscopy. Bioinformatics 31, 1325–1327, doi:10.1093/bioinformatics/btu830 (2015).

42 Muhandiram, D. R. & Kay, L. E. Gradient-Enhanced Triple-Resonance Three Dimensional NMR Experiments with Improved Sensitivity. Journal of Magnetic Resonance, Series B 103, 203–216, doi:10.1006/jmrb.1994.1032 (1994).

43 Grzesiek, S. & Bax, A. Correlating backbone amide and side chain resonances in larger proteins by multiple relayed triple resonance NMR. Journal of the American Chemical Society 114, 6291–6293, doi:10.1021/ja00042a003 (1992).

44 Kay, L., Keifer, P. & Saarinen, T. Pure absorption gradient enhanced heteronuclear single quantum correlation spectroscopy with improved sensitivity. Journal of the American Chemical Society 114, 10663–10665, doi:10.1021/ja00052a088 (1992).

45 Weigelt, J. Single Scan, Sensitivity- and Gradient-Enhanced TROSY for Multidimensional NMR Experiments. Journal of the American Chemical Society 120, 10778–10779, doi:10.1021/ja982649y (1998).

46 Zhang, O., Kay, L. E., Olivier, J. P. & Forman-Kay, J. D. Backbone 1H and 15N resonance assignments of the N-terminal SH3 domain of drk in folded and unfolded states using enhanced-sensitivity pulsed field gradient NMR techniques. J Biomol Nmr 4, 845–858 (1994).

47 Ulrich, E. L. et al. BioMagResBank. Nucleic Acids Res 36, D402–408, doi:10.1093/nar/gkm957 (2008).

48 Ahlner, A., Carlsson, M., Jonsson, B. H. & Lundstrom, P. PINT: a software for integration of peak volumes and extraction of relaxation rates. J Biomol Nmr 56, 191–202, doi:10.1007/s10858-013-9737-7 (2013).

49 Mulder, F. A., Schipper, D., Bott, R. & Boelens, R. Altered flexibility in the substrate binding site of related native and engineered high-alkaline Bacillus subtilisins. J Mol Biol 292, 111–123, doi:10.1006/jmbi.1999.3034 (1999).

50 Mathematica (Wolfram Research, Inc., Champaign, Illinois, 2017).

51 Loria, J. P., Rance, M. & Palmer, A. G. A Relaxation-Compensated Carr-Purcell-Meiboom-Gill Sequence for Characterizing Chemical Exchange by NMR Spectroscopy. Journal of the American Chemical Society 121, 2331–2332, doi:10.1021/ja983961a (1999).

52 Tollinger, M., Skrynnikov, N. R., Mulder, F. A., Forman-Kay J. D. & Kay, L. E. Slow dynamics in folded and unfolded states of an SH3 domain. J Am Chem Soc 123, 1134111352, doi:10.1021/ja011300z (2001).

53 Mulder, F. A. A., Skrynnikov, N. R., Hon, B., Dahlquist, F. W. & Kay, L. E. Measurement of Slow (μs-ms) Time Scale Dynamics in Protein Side Chains by15N Relaxation Dispersion NMR Spectroscopy: Application to Asn and Gln Residues in a Cavity Mutant of T4 Lysozyme. Journal of the American Chemical Society 123, 967–975, doi:10.1021/ja003447g (2001).

54 Mulder, F. A., Hon, B., Mittermaier, A., Dahlquist, F. W. & Kay, L. E. Slow internal dynamics in proteins: application of NMR relaxation dispersion spectroscopy to methyl groups in a cavity mutant of T4 lysozyme. J Am Chem Soc 124, 1443–1451, doi:10.1021/jp0119806 (2002).

55 Guenneugues, M., Berthault, P. & Desvaux, H. A method for determining B1 field inhomogeneity. Are the biases assumed in heteronuclear relaxation experiments usually underestimated? J Magn Reson 136, 118–126, doi:10.1006/jmre.1998.1590 (1999).

56 Mcconnell, H. M. Reaction Rates by Nuclear Magnetic Resonance. J Chem Phys 28, 430–431, doi:Doi 10.1063/1.1744152 (1958).

57 Sekhar, A. et al. Thermal fluctuations of immature SOD1 lead to separate folding and misfolding pathways. Elife 4, e07296, doi:10.7554/eLife.07296 (2015).

58 Tamiola, K., Acar, B. & Mulder, F. A. Sequence-specific random coil chemical shifts of intrinsically disordered proteins. J Am Chem Soc 132, 18000–18003, doi:10.1021/ja105656t (2010).

